# Commensal oral *Rothia mucilaginosa* produces enterobactin – a metal chelating siderophore

**DOI:** 10.1101/2020.02.20.956391

**Authors:** Carla Uranga, Pablo Arroyo, Brendan M. Duggan, William H. Gerwick, Anna Edlund

## Abstract

Next-generation sequencing studies of saliva and dental plaque from subjects in both healthy and diseased states have identified bacteria belonging to the *Rothia* genus as ubiquitous members of the oral microbiota. To gain a deeper understanding of molecular mechanisms underlying the chemical ecology of this unexplored group, we applied a genome mining approach that targets functionally important biosynthetic gene clusters (BGCs). All 45 genomes that were mined, representing *Rothia mucilaginosa, R. dentocariosa* and *R. aeria*, harbored a catechol-siderophore-like BGC. To explore siderophore production further we grew the previously characterized *R. mucilaginosa* ATCC 25296 in liquid cultures, amended with glycerol, which led to the identification of the archetype siderophore enterobactin by using tandem Liquid Chromatography Mass Spectrometry (LC/MS/MS), High Performance Liquid Chromatography (HPLC), and Nuclear Magnetic Resonance (NMR) spectroscopy. Normally attributed to pathogenic gut bacteria, *R. mucilaginosa* is the first commensal oral bacterium found to produce enterobactin. Co-cultivation studies including *R. mucilaginosa* or purified enterobactin revealed that enterobactin reduced growth of certain strains of cariogenic *Streptococcus mutans* and pathogenic strains of *Staphylococcus aureus*. Commensal oral bacteria were either unaffected by, reduced in growth, or induced to grow adjacent to enterobactin producing *R. mucilaginosa* or the pure compound. Taken together with *Rothia*’s known capacity to ferment a variety of carbohydrates and amino acids, our findings of enterobactin production adds an additional level of explanation to *R. mucilaginosa*’s colonization success of the oral cavity. Enterobactin is the strongest Fe(III)-binding siderophore known, and its role in oral health requires further investigation.

**Importance:** The communication language of the human oral microbiota is vastly underexplored. However, a few studies have shown that specialized small molecules encoded by BGCs have critical roles such as in colonization resistance against pathogens and quorum sensing. Here, by using a genome mining approach in combination with compound screening of growth cultures, we identified that the commensal oral community member *mucilaginosa* harbors a catecholate-siderophore BGC, which is responsible for the biosynthesis of enterobactin. The iron-scavenging role of enterobactin is known to have positive effects on the host’s iron pool and negative effects on host immune function, however its role in oral health remains unexplored. *R. mucilaginosa* was previously identified as an abundant community member in cystic fibrosis, where bacterial iron cycling plays a major role in virulence development. With respect to iron’s broad biological importance, iron-chelating enterobactin may explain *R. mucilaginosa*’s colonization success in both health and disease.

## Introduction

Over the past few decades of oral microbiology research, we have come to understand that oral microbiota is imperative not only for our oral health but also for our overall wellness. Thus far, the oral microbiology research field has focused significant efforts on either describing taxonomic shifts of complex bacterial communities in healthy and disease states, or studies of a few model organisms, e.g. cariogenic *Streptococcus mutans* and the periodontal pathogen *Porphyromonas gingivalis*, while less attention has been paid to commensal bacteria such as members belonging to the *Rothia* genus. Our knowledge of bacterial taxonomic signatures expands well beyond our knowledge of the functional roles of oral bacteria. However, some studies of small molecules (SMs) (a.k.a. secondary metabolites), secreted by members of the *Streptococcus* genus, have illustrated that antimicrobial bacteriocins and nonribosomal peptides play crucial roles in colonization resistance against pathogens (1–3). In addition, recent genome mining studies show that the broader oral microbiome is exceptionally rich in biosynthetic gene clusters (BGCs), of which most are unexplored (4–6). Siderophores represent a particularly interesting class of SMs since they not only have the capacity to modulate the human microbiota, but also play critical biological roles for the eukaryotic host by serving either as virulence mediators of pathogens, or as a stabilizer of the human iron pool (7,8). Bacteria release siderophores into the surrounding environment where their apo-form binds metals, resulting in the metal-ligated siderophore, which is imported back into the cell to serve multiple functions in a variety of biochemical mechanisms, such as oxido-reduction of heme-containing proteins (9). Siderophores are also known to reduce oxidative stress deriving from reactive oxygen species (ROS) produced by neighboring bacterial species or human cells (10–12).

Recently, our research team found that the oral microbiome harbors siderophore-like BGCs, of which some belong to the nonribosomal peptide synthetase (NRPS) class (4). Although their structures were not experimentally characterized in this previous study, the NRPS small-molecule products were putatively annotated as griseobactin-like (4). The griseobactin-like BGCs were harbored by oral community members belonging to the ubiquitous and commensal *Rothia* genus, which are previously known for their ability to reduce nitrate to nitrite and thought to limit the growth of aciduric and acidogenic caries pathogens (13,14). *R. mucilaginosa* is one of the most well-studied *Rothia* species and has also been identified as one of the most common community members in healthy subjects as compared to subjects with caries disease (15), primary sclerosing cholangitis (a liver and gall bladder disease) (16), oral squamous cell carcinoma (17), and bronchiectasis (18). In contrast to their beneficial associations, recent studies have also shown that *R. mucilaginosa* is associated with caries disease (19,20), and lung diseases such as pneumonia and cystic fibrosis (CF) (21,22). In addition, a study of the CF microbiota reported that *R. mucilaginosa* boosts the growth of pathogenic *Pseudomonas aeruginosa* via cross-feeding mechanisms (14,22). The functional role of *Rothia* in the human microbiota is not known, but its versatile metabolic capacities such as fermentation of both complex carbohydrates and peptides/proteins into primary metabolites (e.g. free amino acids, short chain fatty acids), suggests that one of its ecological roles may be to provide growth substrates for more specialized community members, such as Gram-negative non-mucous degrading bacteria (i.e. *Pseudomonas aeruginosa* (22)). In this study, we explored further the metabolic capacity of *R. mucilaginosa* ATCC 25296 and show that its genome harbors a catechol siderophore BGC that encodes the archetypal siderophore ‘enterobactin’, and not griseobactin as we predicted in our previous genome mining study (4). Similar enterobactin-BGCs were identified in all of the genomes of two additional *Rothia* species; *R. dentocariosa* and *R. aeria*. Here, we show that pure enterobactin reduced growth of some cariogenic strains of *S. mutans*, a few commensal oral *Streptococcus* species and oral *Actinomyces timonensis*. It also inhibited formation of the yellow pigment staphyloxanthin and growth of methicillin-resistant strains of *Staphylococcus aureus* (MRSA). *S. aureus* is not currently considered a pathogen in oral disease, however, this view is changing as more cases of MRSA and MSSA have been reported in infections of the oral cavity (23–25). Our study also illustrates that enterobactin chelates both zinc and magnesium ions but with less affinity than Fe(III). Conclusively, we describe a new key ecological mechanism that involves the metal-chelating siderophore enterobactin, which not only supports survival and growth of the underexplored yet frequently occurring oral *R. mucilaginosa*, but also reduces growth of both cariogenic and a multi drug-resistant pathogens.

## Results and Discussion

More than 2000 oral bacterial, archaeal and fungal species have been identified to date, and these are known collectively as the human oral microbiota (26). Some bacterial members of the oral microbiota can cause diseases such as dental caries and periodontal diseases, while others provide colonization resistance against pathogens (27–29). The advent of high-throughput sequencing technologies has greatly increased our understanding of the complexity of the oral microbiota and shed new light on not only the diversity of bacterial taxa but also on their biosynthetic potential, i.e. the widely distributed biosynthetic gene clusters (BGCs) that encode small molecules with specialized functions (4–6, 27). Previously, our research team conducted a genome mining survey, targeting BGCs in 461 oral bacterial genomes, which identified a vast unexplored repertoire of ∼5,000 putative BGCs (4). In another study of children with deep dentin caries disease, we found that bacterial community members belonging to the *Rothia* genus, specifically *Rothia mucilaginosa*, were enriched and actively replicating DNA in saliva from healthy children as compared to saliva from children with caries (15). From the same metagenomes we also identified BGCs that were identical to those harbored by the complete genome sequence representing *Rothia mucilaginosa* ATCC 25296 (4). Based on this we employed the oral *Rothia mucilaginosa* ATCC 25296 as a model species in this study. Here we expanded on our previous findings and conducted BGC mining of additional *Rothia* genomes and draft genomes, available in NCBI GenBank, to explore to what extent species within this genus harbor unique BGC signatures, which could explain some of *Rothia’*s competitive success in the oral cavity. By using the antiSMASH 5.0 software (bacterial version) (30) we predicted that each of the 26 *Rothia mucilaginosa* genomes and draft genomes (available as of February 2, 2020 at https://www.ncbi.nlm.nih.gov/genome/genomes/1812) harbored a NRPS catechol siderophore-like (cat-sid) BGC (Fig. S1). Each of the 12 *R. dentocariosa* draft genomes (available at https://www.ncbi.nlm.nih.gov/genome/genomes/1968) harbored a butyrolactone BGC, a T1PKS BGC, a lanthipeptide BGC, and a cat-sid BGC. Furthermore, each of the seven *R. aeria* genomes (available at https://www.ncbi.nlm.nih.gov/genome/genomes/12163) harbored a cat-sid BGC and a lasso peptide-like BGC. Taken together, by analyzing genomes representing three different *Rothia* species, unique BGC signatures were observed at the species level and a highly similar NRPS-encoded cat-sid BGC was shared between all three species (Fig. S2), which illustrates a broader ecological importance of this siderophore. AntiSMASH predicted that the closest homologue to the cat-sid BGC was a mirubactin BGC (14% peptide sequence similarity) (Fig. S1C). Mirubactin is a mixed catecholate and hydroxamate-type NRPS siderophore. However, AntiSMASH also predicted that the core building blocks for siderophore biosynthesis were serine and dihydroxy benzoic acid, showing support for the biosynthesis of a catecholate siderophore (Fig. S1).

### Screening for the siderophore in *Rothia mucilaginosa* growth extracts

To test if *Rothia mucilaginosa* ATCC 25296 secreted a siderophore while growing in liquid growth cultures, we tested growth in multiple conditions. We specifically optimized for growth in minimal media to reduce the number of metabolites that could interfere with our downstream mass spectrometric analysis and purification of the siderophore. Plateauing growth curves of *R. mucilaginosa* ATCC 25296 were obtained in liquid minimal medium M9 cultures supplemented either with 100 mM sucrose or glucose during aerobic incubation in 37° C (Fig. 1A). However, when incubated with other carbon sources (glycerol, lactose, arabinose, or galactose) growth was reduced (Fig. 1A). To explore if a siderophore was produced in any of the above culture conditions, we screened liquid growth extracts using two different assays; a hydroxamate assay that targets carboxylate siderophores (31) and the Arnow’s assay, which targets catecholate siderophores (32). Arnow’s assay showed clear colorimetric changes as the normally colorless M9 media turned ruby red in cultures incubated with sucrose and glycerol, while the hydroxamate assay showed no color change. The presence of a catecholate siderophore in these cultures was also confirmed using absorbance measurements at 500 nm, which is known to capture catecholate derivatives (33). Not only did these results illustrate that *R. mucilaginosa* ATCC 25296 can produce a catechol siderophore but also that the AntiSMASH program could accurately predict the correct core building blocks (serine and dihydroxy benzoic acid). To facilitate compound isolation and purification, we further explored if glycerol, which is known to elicit secondary metabolite production in other Actinobacteria (34), could increase siderophore yields. This was indeed the case as enterobactin production increased significantly in the glycerol amended cultures (Absorbance at 500nm: ∼0.25) (Fig 1B). However, under the same conditions, growth decreased (Fig. 1A). We also tested if liquid cultures of *Rothia dentocariosa* M567, which also harbors a cat-sid BGC (Figure S2), can produce a siderophore when subjected to glycerol cultivation (Fig. 1B). Production was observed for this species as well (Absorbance at 500nm: ∼0.15) but not to the same extent as for *R. mucilaginosa* (Fig.1B).

**Figure 1.**
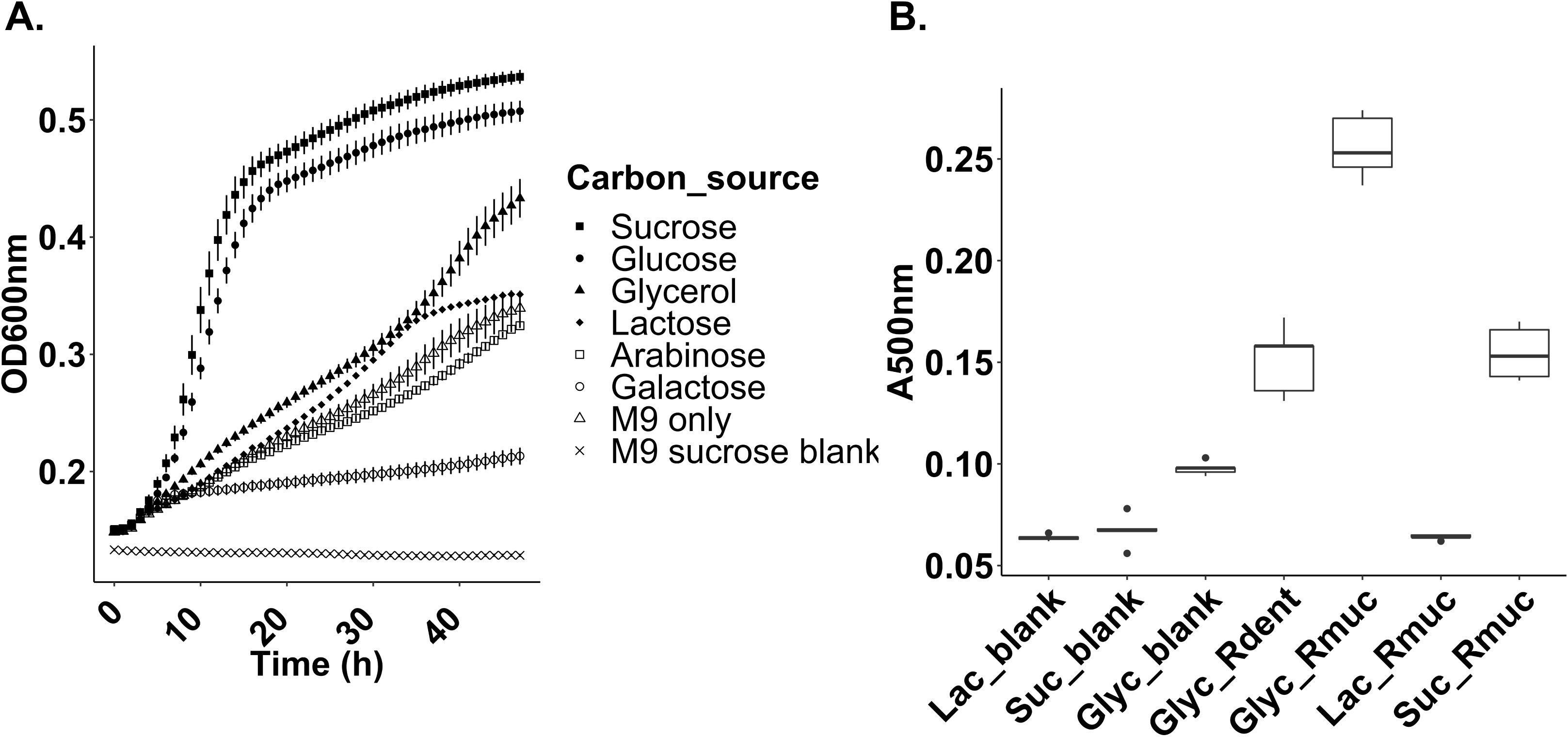
A) Growth curves for *Rothia mucilaginosa* ATCC 25296 (Rmuc) incubated in aerobic conditions in liquid M9 media, supplemented with different carbon sources (x-axis: hours, y-axis: optical density [OD600nm]). B) Colorimetric absorbance at 500 nm (y-axis: A500nm) capturing catecholate derivatives in liquid Rmuc and *R. dentocariosa* M567 (Rdent) growth cultures (x-axis) using Arnow’s assay. Rmuc was grown in M9 media supplemented with 100 mM glycerol (Glyc_Rmuc), 100 mM lactate (Lac_Rmuc), 100 mM sucrose (Suc_Rmuc). Rdent was grown in 100 mM glycerol to see if glycerol induced siderophore production as seen for Rmuc cultures.

### Detailed characterization of *R. mucilaginosa*’s catechol siderophore and cat-sid BGC

Using Phyre2, a protein structure prediction software (35), only one gene within the enterobactin BGC from *R. mucilaginosa* ATCC 25296 showed amino acid sequence homology to the BGC that encodes enterobactin in the *E. coli* JM109 genome (i.e. amino acid location 51-463 in the NRPS gene, Fig. S4). The overall low structural BGC homology as well as the low amino acid sequence homology between the NRPS genes in *R. mucilaginosa* and *E. coli* BGCs illustrate that the local gene environments surrounding the core biosynthetic NRPS genes have diversified greatly between different taxonomic groups of bacteria and have no influence on the final compound structure.

Preliminary high-resolution mass spectrometry (LC-MS/MS) analysis of a *R. mucilaginosa* ATCC 25296 siderophore-enriched sample (derived from a glycerol amended growth culture) in negative ionization mode yielded a compound with a parent mass [M+H] of 670.152 Da. Upon collision induced dissociation (CID) fragmentation of the parent mass, and comparative analysis with all previously characterized compound spectra available through the Global Natural Products Social Molecular Networking (GNPS) library (36), it became clear that the ion fragments matched the well-characterized catecholate siderophore, enterobactin (*m/z* 670.164, GNPS gold standard spectrum CCMSLIB00005435752, GNPS cosine score of 0.71-0.78, Fig. 2A). To verify our findings, we performed additional LC-MS/MS on purified extracts from thin layer chromatography, which again resulted in the identification of enterobactin or a close homolog using the GNPS infrastructure, this time in positive ionization mode (Fig. 2B). Further purification of this compound using HPLC showed a well-separated peak that tested positive in the Arnow’s assay (Fig. 3). This peak was eluted and analyzed by ^1^H NMR to confirm that *R. mucilaginosa* ATCC 25296 produces enterobactin (Table S1). All observed chemical shift values in Table S1 (obtained from *R. mucilaginosa’s* enterobactin) were also reported previously in an Infrared (IR) and NMR spectroscopic study of the archetypal enterobactin molecule (40). Proton and 2D NMR spectra (HSQC, HMBC, and COSY) for the *R. mucilaginosa*-derived compound confirmed its identity (Figure S5).

**Figure 2.**
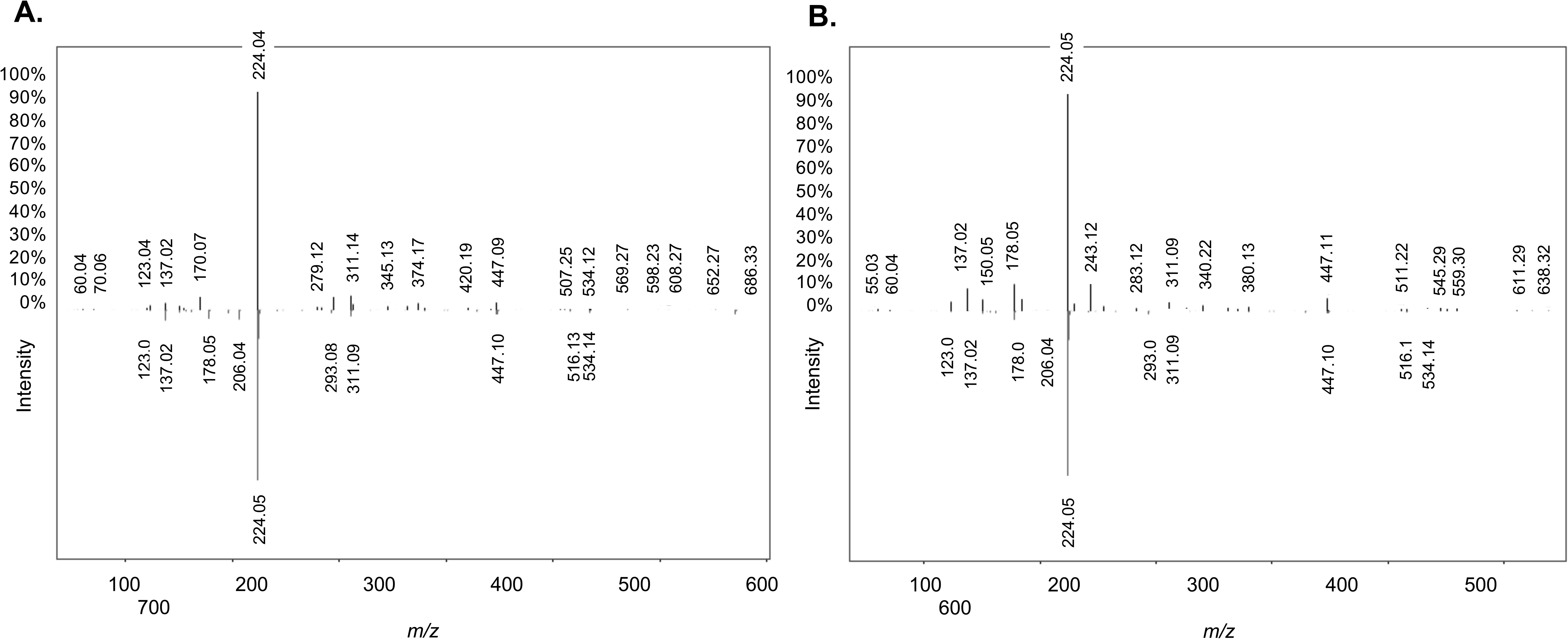
Replicate mirror mass fragmentation patterns for enterobactin (*m/z* 670.152) produced by *Rothia mucilaginosa* ATCC 25296 (ion fragments pointing up) and enterobactin from the gold standard spectrum (ion fragments pointing down) in the Global Natural Products Social Molecular networking (GNPS) library (39). Six major fragments (*m/z* 123.04, *m/z* 137.02, *m/z* 224.05, *m/z* 311.09 – 311.14, *m/z* 447.09 - 447.11, *m/z* 534.14) of the query compound matched the gold standard in GNPS. A) Purified extract derived from *R. mucilaginosa* ATCC 25296 growth media (negative ionization mode). B) Further enrichment of the siderophore using thin layer chromatography (positive ionization mode). Both experiments confirmed the production of enterobactin.

**Figure 3.**
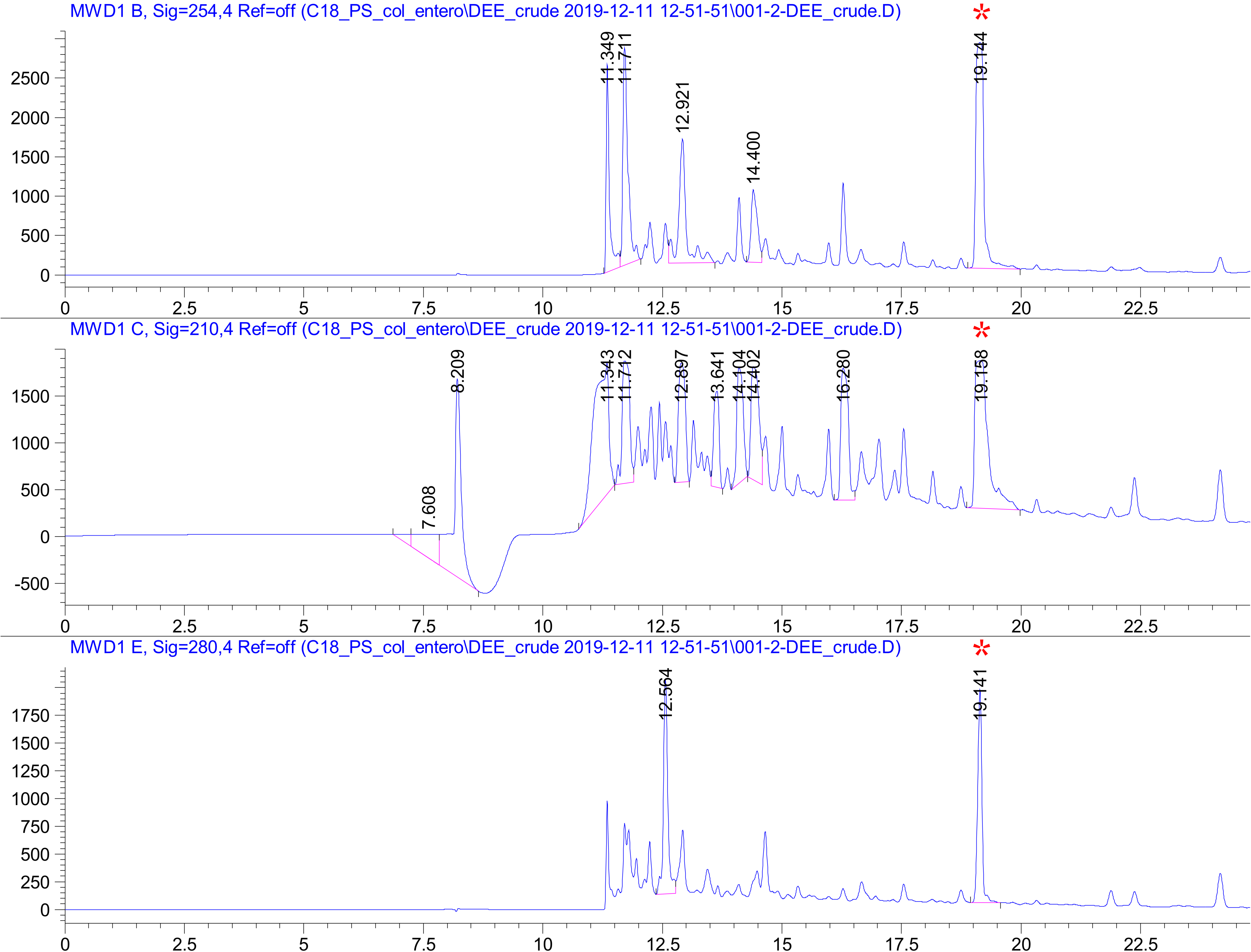
High Performance Liquid Chromatography (HPLC) purification traces of enterobactin from *Rothia mucilaginosa* ATCC 25296 crude growth extract, measured at 210 nm, 254 nm and 280 nm using a solvent gradient from 30-65% buffer B. The peak at 19.141 min (red asterisk) was eluted and further analyzed by NMR. X-axis: minutes. Y-axis: peak absorbance intensity.

### Enterobactin activity screening using co-cultivation assays

Interactions between *R. mucilaginosa* ATCC 25296 or the purified enterobactin compound and other bacterial species were studied by employing co-cultivation agar assays and liquid growth assays. On agar plates, growth of *R. mucilaginosa* was established prior to spotting the challenging species. Interactions were initially studied on both rich (BHI) and minimal M9 agar media to investigate under which conditions *R. mucilaginosa* could modulate growth of the competitor test strain, potentially via iron competition (visualized as clearing zones surrounding *R. mucilaginosa*). In the liquid mono-cultures, the purified siderophore was added at the same time as the cultures were seeded with the test strain. We found that while growing on minimal M9 agar, supplemented with glycerol (100 mM) during plate interaction studies, *R. mucilaginosa* inhibited growth of *Actinomyces timonensis* DSM 23838 (Fig. S3A), and reduced pigment production in four strains of pathogenic *Staphylococcus aureus* in the presence of catalase (Fig. 4, Fig. S6). We added catalase to the agar plates to prevent growth inhibition by eventual reactive oxygen species (ROS) produced by *R. mucilaginosa*. The results suggest that inhibition of pigment production is not due to ROS but eventually to enterobactin. To further elucidate this interaction, we conducted mono-culture experiments on agar plates where we added the purified enterobactin to *S. aureus* agar plates (Fig. 5). With catalase added, statistically significant differences in pigmentation were observed for all *S. aureus* strains amended with enterobactin, including MRSA strains TCH70/MRSA and NR10129 (p<0.05, two-tailed t-test) (Fig. 5), confirming an important role of enterobactin in the inhibition of *S. aureus* virulence and growth (37,38). For all the tested oral commensal and pathogenic *Streptococcus* species, no growth inhibition was observed on agar plates. However, changes in growth were confirmed for some of the species when adding the purified enterobactin compound to liquid cultures (with or without catalase) (Fig. 6). Of the pathogenic *Streptococcus* strains tested in liquid growth cultures amended with enterobactin and catalase, we observed that growth of both *S. mutans* UA159 and *S. mutans* B04Sm5 were significantly reduced (p<0.05, t-test of averaged mean values) (Fig. 6A, 6B), illustrating that *R. mucilaginosa* has the capacity to reduce growth of cariogenic pathogens, potentially via iron competition. The commensal oral bacteria that were tested showed varying responses to enterobactin and catalase (for a list of species tested see Materials and Methods section). For example, growth of *S. sanguinis* ATCC 49296 was not significantly impacted by enterobactin (Fig. 6C). However, when subjected to catalase treatment (with and without enterobactin), *S. sanguinis* growth rate increased as optical density (OD600) reached 0.25 after 10 hours of growth as compared to the non-catalase amended cultures (OD 0.25 was reached after 20 hours) (Fig. 6C). This reflects that catalase protects *S. sanguinis* from oxidative damage caused by its own ROS production and thereby stimulates its growth (39). Growth of *S. gordonii* ATCC 35105 was significantly enriched by enterobactin but only without catalase (p<0.0001, t-test of averaged mean values), which illustrates that growth was boosted by the presence of enterobactin (Fig. 6D). A similar trend was observed for *Streptococcus salivarius* SHI-3, which also increased significantly in cultures amended with enterobactin when no catalase was added (p<0.05, t-test of averaged mean values) (Fig. 6E). Conversely, *S. oralis* grew significantly more in the presence of catalase only (p<0.0001, t-test of averaged mean values), while growth was reduced by enterobactin (p< 0.05, t-test of averaged mean values) (Fig. 6F). These results show that bacteria belonging to the *Streptococcus* genus have a varied response to enterobactin in the presence or absence of ROS. The liquid culture experiments illustrate that *S. salivarius* and *S. gordonii* can benefit from the presence of enterobactin-producing *R. mucilaginosa* while *oralis* growth can be reduced. A growth boost of *S. salivarius* was also observed on co-culture agar plates (minimal M9 media supplemented with glycerol) adjacent to *R. mucilaginosa* (Figure S3B). Whether this response was due to the presence of enterobactin needs to be explored further.

**Figure 4.**
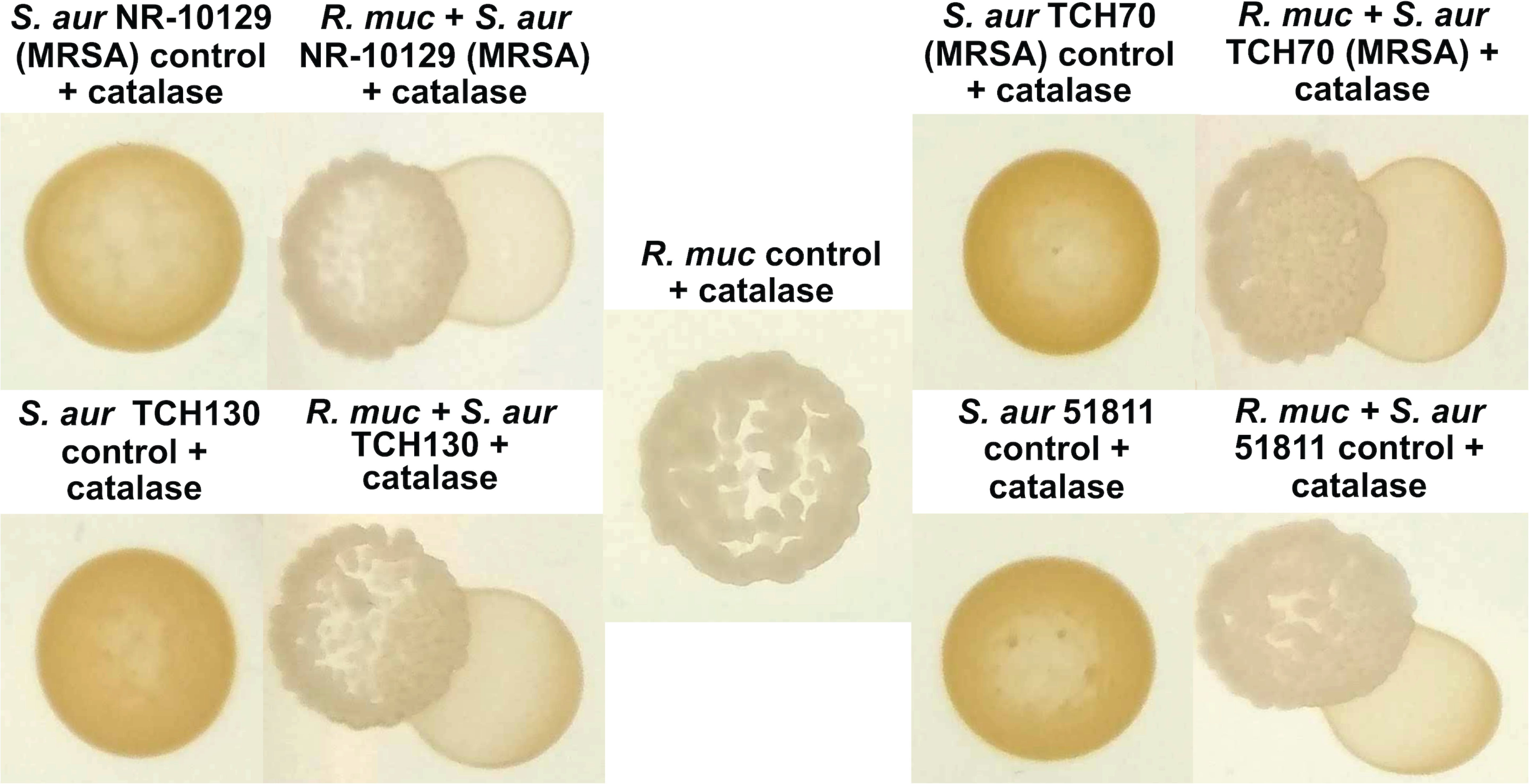
*Rothia mucilaginosa* ATCC 25296 (R. muc) inhibits pigment production in *Staphylococcus aureus* (MRSA) NR-10129, *S. aureus* TCH70 (MRSA), *S. aureus* TCH130 and enterotoxin H producing *S. aureus* ATCC 51811, on M9 agar plates (100 mM glycerol) with 8 µg/mL catalase added. All *S. aureus* strains show yellow pigmentation when growing alone.

**Figure 5.**
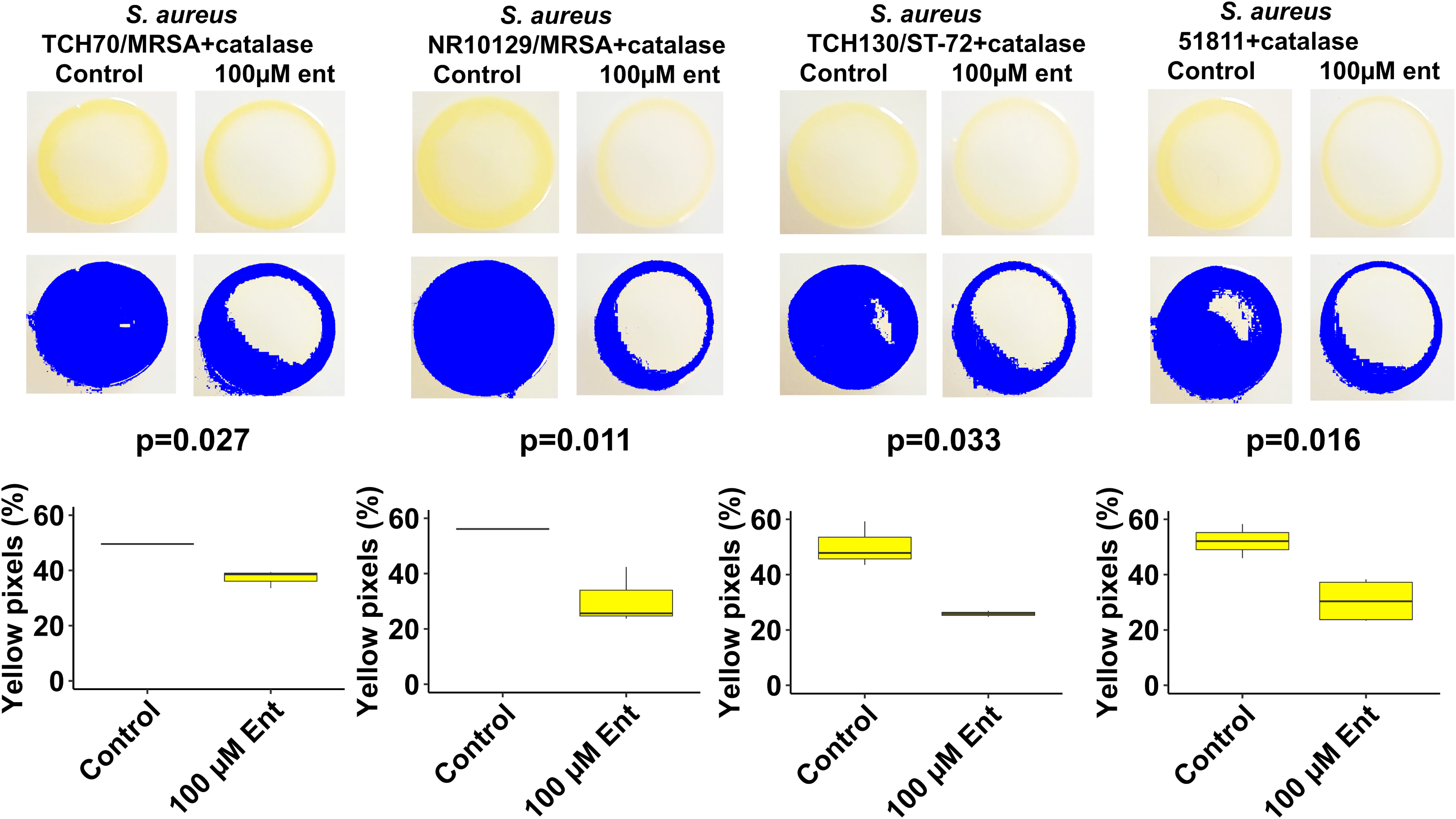
Yellow pigmentation of *S. aureus* strains exposed to 100 µM enterobactin and 8 µg/mL catalase on M9 agar plates supplemented with100 µM glucose. Yellow pigmentation was measured with the R package “countcolors” (51). All strains presented statistically significant reduction in pigmentation in the presence of 100 µM enterobactin purified from *R. mucilaginosa* ATCC 25296 and catalase (p<0.05, two-tailed t-test). Boxplots from yellow pixel measurements were generated with the R Studio program (49) and ggplot2 (43).

**Figure 6.**
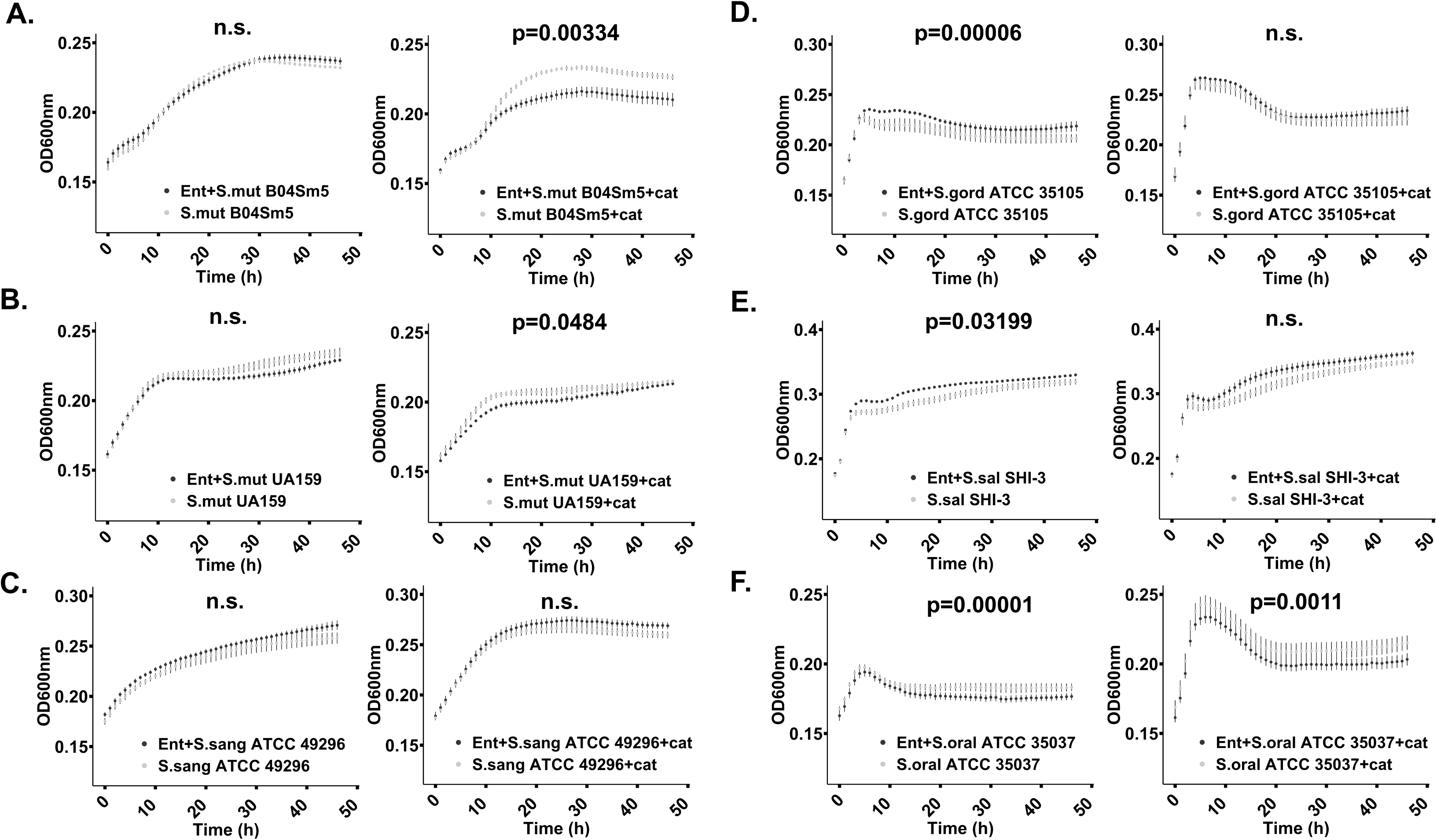
Growth curves of cariogenic and commensal *Streptococcus* species. Bacteria were grown aerobically in 37°C in liquid M9 media supplemented with 100 mM glucose, 1 µM FeCl_3_, either with or without 100 µM enterobactin purified from *R. mucilaginosa*, with or without 8 µg/mL catalase. Growth curves in black represent cultures amended with enterobactin. Statistically significant growth curves (p<0.05) are shown with corresponding p-values. Error bars reflect the standard error of the mean (calculated from triplicates). A) Growth of cariogenic *S. mutans* strain B04Sm5 with enterobactin only (left panel), and with enterobactin and catalase (right panel). B) Growth of cariogenic *S. mutans* strain UA159 with enterobactin only (left panel), and with enterobactin and catalase (right panel). C) Growth of commensal *S. sanguinis* ATCC 49296 with enterobactin only (left panel), and with enterobactin and catalase (right panel). D) Growth of *S. gordonii* ATCC 35101 with enterobactin only (left panel), and with enterobactin and catalase (right panel). E) *S. salivarius* strain SHI-3 with enterobactin only (left panel), and with enterobactin and catalase (right panel). F) Growth of *S. oralis* ATCC 35037 with enterobactin only (left panel), and with enterobactin and catalase (right panel). Graphs were generated and statistically validated using the R studio (41) and the “statmod” and “ggplot2” packages (48,50).

Liquid growth culture experiments including enterobactin were also performed for pathogenic *Staphylococcus aureus* strains. Growth of the enterotoxin H producing strain *S. aureus* ATCC 51811 and the MRSA strain *S. aureus* TCH130 130/ST-72 were significantly reduced by enterobactin when catalase was present, which suggests that enterobactin negatively impacts *S. aureus* growth (p<0.0005 and p<0.005, t-test of averaged mean values, respectively) (Figs. 7A, 7B). None of the other tested MRSA strains showed reduced growth in enterobactin amended cultures (with or without catalase added) (Figs. 7C, 7D). Interestingly, our findings illustrate that strains belonging to the same bacterial species show different responses to enterobactin, which warrant further investigations of the role of strain level sensitivity to this compound in *S. aureus* MRSA virulence and growth.

**Figure 7.**
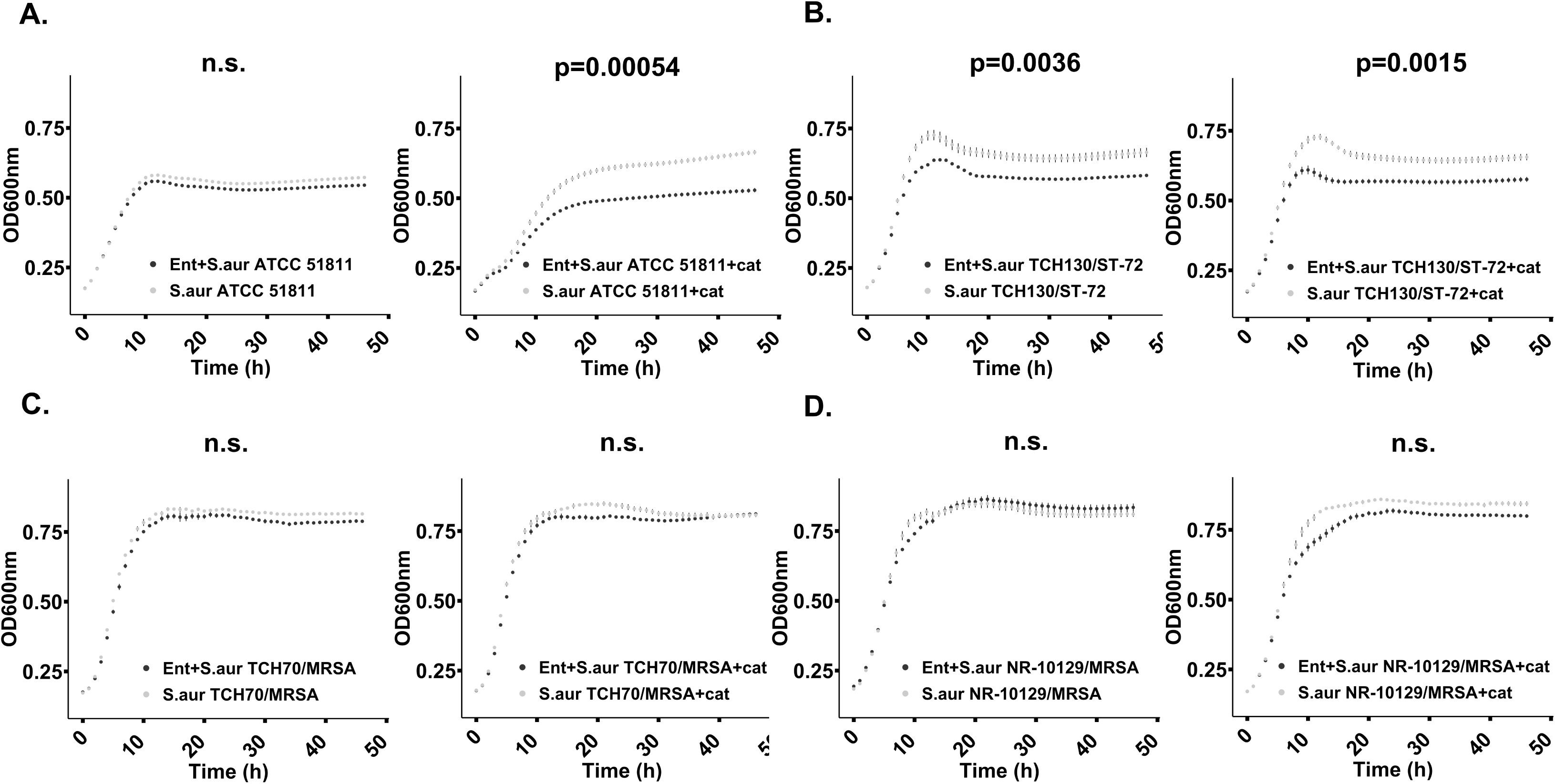
Reduced growth of *Staphylococcus aureus* incubated in liquid M9 growth media (100 µM enterobactin, 8 µg/mL catalase, 100mM glucose) at 37°C for 24 hours. No enterobactin was added to the control samples. Growth curves in black represent cultures amended with enterobactin. Statistically significant growth curves (p<0.05) are shown with corresponding p-values. Error bars reflect the standard error of the mean (calculated from triplicates). A) Growth of *S. aureus* strain 51811 with enterobactin only (left panel) and with enterobactin and catalase (right panel). B) Growth of *S. aureus* strain TCH130/ST-72 with enterobactin only (left panel) and with enterobactin and catalase (right panel). C) Growth of *S. aureus* TCH70/MRSA with enterobactin only (left panel) and with enterobactin and catalase (right panel). D) Growth of *S. aureus* NR-10129/MRSA with enterobactin only (left panel) and with enterobactin and catalase (right panel). Graphs were generated and statistically validated using the R studio (41) and the “statmod” and “ggplot2” packages (48,50).

To evaluate if enterobactin was actively chelating free iron throughout the liquid cultivation experiments, we incubated 500 µM of the purified compound in sterile M9 liquid media as described for bacterial liquid cultures. By using Arnow’s assay we observed that its binding capacity to molybdenum (which in Arnow’s assay substitutes for iron) was reduced by 14% after six hours of incubation. Assuming that this response is linear we estimated that 56% of enterobactin’s activity was lost after 24 hours. This suggests that the compound became inactive in our co-culture experiments over time for physiochemical reasons and that a more dramatic effect of enterobactin likely would have been observed if its activity had remained stable.

We also investigated if enterobactin can bind other metal ions by using the calmagite assay, which is normally used for testing the hardness of water by detecting ions such as magnesium (Mg), calcium (Ca) and zinc (Zn) that bind to ethylenediaminetetraacetic acid (EDTA) (40,41). EDTA is used as a reference compound due to its well-known chelation of divalent cations. Results from this test revealed that enterobactin can chelate both Mg^2+^ and Zn^2+^ ions. Enterobactin showed a chelation activity of 40 µM EDTA-equivalents at a concentration of 100 µM, which is 2.5 times less than EDTA affinity for Mg^2+^ and Zn^2+^ (Fig. S7). These results illustrate that enterobactin can indeed chelate additional ions and not only ferrous iron (Fe^3+^), which could have critical impacts on other microbial community members as well as the human host. It is well known that Zn^2+^ is a critical component of the ubiquitous protein zinc fingers that are able to interact directly with DNA, RNA, and proteins. Magnesium (Mg^2+^) is involved in nervous system signaling, immune system function and bone formation. Our findings show that *R. mucilaginosa* has the potential to compete with the host not only for Fe^3+^ but also for Mg^2+^ and Zn^2+^, which suggest that *R. mucilaginosa* could play an important role in health and disease outcomes.

In conclusion, our study shows that *R. mucilaginosa* ATCC 25296 produces the archetype siderophore enterobactin when growth conditions are suboptimal in a minimal growth media supplemented with glycerol; a known inducer of secondary metabolites in *Streptomyces* (42). We observed different growth responses to purified enterobactin by various members of the oral microbiota; some commensal *Streptococcus* species increased in growth while others decreased (independently in the presence of ROS). Our findings also illustrate that growth of pathogenic bacteria (i.e. two cariogenic strains of *S. mutans* and methicillin-resistant strains of *S. aureus*) is reduced in growth in cultures amended with enterobactin. Moreover, while methicillin-resistant *S. aureus* grew adjacent to enterobactin producing *R. mucilaginosa* ATCC 25296 (with or without ROS present), the virulence factor known as golden pigmentation (staphyloxanthin) was highly reduced. While we provide evidence that the loss of this golden pigmentation is caused by enterobactin, the molecular mechanisms of the interaction remain to be explored. Further examination of the role of *R. mucilaginosa* enterobactin in inhibiting pigment production and bacterial growth may provide a pathway towards the development of new therapeutic leads against not only MRSA strains but also other pathogens, such as cariogenic *Streptococcus mutans*.

## Materials and Methods

### Bacterial isolates used in this study

*Rothia mucilaginosa* ATCC 25296, *Rothia dentocariosa* M567, *Streptococcus mitis ATCC 6249, Streptococcus salivarius* SHI-3 (isolate from oral *in vitro* biofilms) (43), *Streptococcus sanguinis* ATCC 49296, *Streptococcus oralis* ATCC 35037, *Streptococcus gordonii* ATCC 35105, *Actinomyces timonensis* DSM 23838, *Streptococcus mutans* UA159, *Streptococcus mutans* B04Sm5, *Staphylococcus aureus* strain MN8 (MSSA), *Staphylococcus* aureus strain NR-10129 (MRSA), *Staphylococcus aureus* strain TCH130/ST-72, *Staphylococcus aureus* ATCC 51811 (enterotoxin H producer), *Staphylococcus aureus* TCH 70 (MRSA).

### Growth media used in this study

A minimal medium was modified and developed from the M9 media recipe reported by Ebling and Brent (44). All glassware was rinsed thoroughly with 2.5 M HCl and washed with DI H_2_O before use. The fundamental M9 media formulation (pH 7) was supplemented with 0.8% deferrated acid-hydrolyzed casamino acids (BD Biosciences San Jose, CA, USA), 8 mM MgSO_4,_ (Millipore-Sigma) and with various carbon sources (sucrose, glycerol, galactose, lactose, arabinose, glucose or lactate), to a final concentration of 100 mM. The acid-hydrolyzed casamino acids were deferrated with an equal volume of 3% 8-hydroxyquinoline (Fisher Chemical, Pittsburgh, PA, USA) in chloroform. MgSO_4_, glucose and sucrose were added post-autoclaving to prevent solution clouding and the Maillard reaction (“caramelization” of sugars with amino acids). Plates were made of the same formulations, containing 1% agar. Nutrient and iron-rich Brain heart infusion (BHI) media (Oxoid, Thermo Scientific, Carlsbad, CA, USA) were also used in liquid cultures and agar plates for cultivating experimental strains.

### Mining for biosynthetic gene clusters (BGCs) in Rothia genomes

The BGC prediction program antiSMASH, bacterial version (30), which is able to predict core secondary metabolite structures from BGC sequences, was used to identify BGCs in 26 genomes (including both completed and draft genomes) representing *R. mucilaginosa*, available at https://www.ncbi.nlm.nih.gov/genome/genomes/1812?, 12 genomes (including both completed and draft genomes) representing *R. dentocariosa*, available at https://www.ncbi.nlm.nih.gov/genome/genomes/1968?, and 7 genomes (one full length and four draft genomes) representing *R. aeria*, available at https://www.ncbi.nlm.nih.gov/genome/genomes/12163?. The NaPDoS program was used for further classification of the non-ribosomal peptide synthase (NRPS) catechol siderophore BGCs by using the C-domain classification tool (45).

### BGC Structural analysis

For structural homology modeling and comparative analysis of the NRPS protein harbored by the *Rothia mucilaginosa* ATCC 25296 cat-sid BGC, Phyre2 software (35) was used to compare the NRPS with known structures in the Phyre2 database.

### Co-cultivation agar assays

For bacterial interaction screening studies, all bacterial isolates were seeded from glycerol stocks in BHI (Oxoid, Thermo Scientific) media and incubated for 24 h before placing on either BHI agar or M9 minimal agar media supplemented with either 100 mM sucrose (Acros Organics, Pittsburgh, PA, USA), glucose (Millipore-Sigma), or glycerol (Honeywell, Mexico City, Mexico). Plates were incubated either in a 5% CO_2_ incubator at 37°C or anaerobically at 37°C in an anaerobic chamber (Coy Laboratory Products, Grass Lake, MI) with a gas mix of 5% H_2_, 5%CO_2,_ and 90% N_2_. All tested strains were grown similarly except *A. timonensis*, which was incubated for 48 h in BHI in an anaerobic chamber at 37°C before plating due to its slower growth rate in M9 minimal media. *R. mucilaginosa* or *R. dentocariosa* were plated by dropping 20-30 µL, in three to six replicates, onto plates, allowing the drops to air-dry before incubation. Control incubations consisted of 20-30 µL drops of each tested bacterial species. Agar plates with bacteria were incubated 24-48 h prior to adding another species. Interactions with other bacteria were tested by adding a 20-30 µL drop next to the first strain, making sure the drop made contact with the established bacterial strain. Interactions were assessed after another 24-76 h of incubation. Interactions were screened with the naked eye for growth inhibitions zones or other growth-related behaviors. All interaction assays were repeated at least three times and documented.

### Siderophore detection assays

To clarify the type of siderophore produced by *R. mucilaginosa* ATCC 25296 *and R. dentocariosa* M567, siderophore-positive supernatants were assayed using Arnow’s assay (32) for catecholate compounds, and the iron perchlorate assay for hydroxamate siderophores (31). For the hydroxamate siderophore assay, 1.0 ml of a 10 mM solution of Fe(CIO_4_)_2_ (Alfa Aesar, Tewksbury, MA) in 0.1M HClO_4_ (Ricca Chemical, Visalia, CA, USA) was mixed with 1.0 ml of unknown and the absorbance read at 495 nm. For Arnow’s assay, cultures were screened by mixing 200 µL culture with 20 µL 5M HCl (Ricca Chemical), followed by 100 µL Arnow’s reagent (20% sodium molybdate dihydrate (EMD St. Louis, MO, USA) and 20% sodium nitrite (Fisher chemical) in DI H_2_O. To develop the ruby red color indicating the presence of a catecholate siderophore, 20 µL 10 N NaOH (Ricca Chemical) was added. These were assessed visually for culture and HPLC fraction screening. For quantitative assays these were measured spectrophotometrically at 500 nm.

### Siderophore purification and structural elucidation

Siderophore enrichment in growth cultures of *R. mucilaginosa* ATCC 25296 was implemented in several steps. First, after 7 days of incubation, the supernatant of a liquid culture was acidified to pH 2. Twenty grams per 100 mL of amberlite XAD16 (Alfa Aesar) was added to the supernatant, which were then placed in an orbital shaker at 175 rpm, and shaken overnight. The amberlite was collected, washed three times with DI H_2_O, then extracted with diethyl ether (DEE) (Millipore-Sigma) and methanol (Millipore-Sigma). The solvent phase (the top diethyl ether layer) was evaporated at room temperature in an open container. The material was separated on silica gel thin layer chromatography (TLC) (Millipore-Sigma) and developed in a mixture of 65% n-butanol (Acros Organics), 25% acetic acid (Fisher Scientific, Pittsburgh, PA, USA), and 15% water, then developed with Arnow’s reagent. The Arnow-positive TLC band was removed and re-extracted with DEE, solvent evaporated and the remaining material tested with Arnow’s assay. Mass spectrometry (LC-MS/MS) using a triple TOF mass spectrometer (AB Sciex 5600, Framingham, MA, USA) was done in both positive and negative modes on a PS C18 column (Phenomenex, Torrance, CA; 2.6 µM particle size, 4.6 mM diameter, 250mM length) and eluted with a gradient of 0-100% acetonitrile, and 0.1% formic acid (Honeywell, Mexico City, Mexico). Upon obtaining data, the msConvert program was used to convert vendor files to the mzXML format (46). The Global Natural Products Social Molecular Networking (GNPS) platform was used to putatively annotate the detected fragment masses (47). For siderophore purification, the crude DEE extract was resuspended in 50% methanol/water and purified with the PS C18 column using a 20-minute gradient of 30%-65% buffer B (acetonitrile with 0.1% formic acid) using an Agilent 1200 series HPLC with a fraction collector. Buffer A consisted of water with 0.1% formic acid. The pure HPLC fraction was diluted with DI H_2_O, frozen at −80°C and lyophilized to remove solvents and water. For NMR, pure lyophilized enterobactin was resuspended in DMSO-d_6_ and analyzed via nuclear magnetic resonance (NMR) with a 600 MHz (14.1 Tesla) Bruker Avance III NMR fitted with a 1.7 mm inverse detection triple resonance cryoprobe with z-gradients.

### Liquid co-cultivation assay with pure enterobactin

In order to gain further insight into the effects of enterobactin on bacterial growth, liquid growth experiments were prepared with purified enterobactin from *R. mucilaginosa*. Because hydrogen peroxide can also be responsible for growth inhibition, catalase (also known as hydrogen peroxide oxidoreductase) was used in the culture medium to degrade hydrogen peroxide and isolate the effect of the siderophore itself. Briefly, to enrich for growth of each bacterial species that were included in co-cultivation experiments in minimal M9 media, cultures were first established in BHI from frozen glycerol stocks and incubated for 24 h aerobically in 37°C with 5% CO_2_. For the growth curve assays, glucose was chosen as a universal carbon source to accommodate all strains used, some of which are not able to grow on glycerol. BHI media was removed from the cells prior to starting experiments in minimal M9 media as follows: one mL of each culture was centrifuged at 7000 rpm for 10 minutes, the supernatant was removed from the pellet using a sterile pipette, and the pellet was re-suspended in an assay media consisting of 0.5 mL minimal M9 media supplemented with 100 mM glucose, 8 mM MgSO_4_, 0.8% deferrated amino acids, 1 µM FeCl_3_, either with or without 8 µg/mL catalase (Millipore-Sigma). The bacterial cell suspensions were further diluted 1:20 in the same media. A stock solution of enterobactin was prepared for the assay by suspending purified enterobactin in M9 + 100 mM glucose, 8 mM MgSO_4_, and 0.8% deferrated amino acids with or without catalase) to 200 µM. One hundred µL of the 200 µM siderophore suspension were added to a 96 well plate, in triplicate, for each bacterial strain studied. One hundred µL of the 1:20 bacterial suspension was added to each well, in triplicate for each condition. The control consisted of the same preparations, but without the siderophore. All plates were incubated at 37°C for 48 h in a Tecan Infinite M Nano spectrophotometer and growth was monitored every hour by absorbance measurements at 600 nm. Statistical analysis was done with the R package “statmod” using the “compareTwoGrowthCurves” function with an nsim parameter value of 100000 (48) based on a meanT calculation. All graphical figures were generated using R Studio, version 1.2.5001 (49) and the package ggplot2 (50).

### Agar plate assays of S. aureus strains amended with pure enterobactin from R. mucilaginosa

To assess growth and staphyloxanthin pigmentation (a.k.a. golden pigment) production of *S. aureus* strains on M9 agar plates supplemented with glucose and pure enterobactin, 24 h cultures of *S. aureus* strains grown in BHI were pre-mixed 1:2 with a 2X solution containing 2X M9 media, 200 mM glucose, 16 µg/mL catalase and 200 µM enterobactin so that final concentrations were 1XM9, 100 mM glucose, 8 µg/mL catalase and 100 µM enterobactin. Controls consisted of the exact same formulation but without enterobactin. Twenty µL drops were plated onto M9 agar plates (100mM glucose) as three replicates and incubated 24 h at 37°C and 5% CO_2_. The R package “countcolors”, version 0.9.1 (51) was used for quantifying the level of yellow pigmentation in the bacterial growth areas by setting a pixel color detection range using only the RGB values detected in the images obtained. A rectangular range of color (RGB scale) values for yellow pigmentation detection used were upper RGB values of 0.988235294, 0.960784314, 0.823529412, and lower RGB values of 0.91372549, 0.850980392, 0.396078431. The fractions of yellow pixels per image were quantified in this way and converted to percentage values, indicating the percentage of yellow pixels detected in each image. The command in the “countcolors” package used was “countcolors::countColorsInDirectory”, in order to automate yellow pigmentation parameters for all images in the image folder and to standardize detection across the strains and replicates. Separate images were created with the results in order to illustrate the detection of yellow pixels. These separate images show the pixels that were detected and counted by the R program “countcolors”. The yellow pixels were automatically replaced with blue by the program to better visualize the effect of enterobactin on *S. aureus* pigmentation. Three images per condition and strain were measured for yellow pigmentation, and a two-tailed T-test performed for each strain using the percentage values obtained by the “countcolors” program, comparing each strain with a control strain as described above.

### Stability testing of enterobactin

To test the stability of enterobactin in conditions used in the liquid growth experiments, pure enterobactin was resuspended at 1 mM in H_2_O and diluted 1:2 to 500 µM in a 2X solution of 2X M9, 200mM glucose, 2 µM FeCl_3_, either with or without 16 µg/mL catalase. Twenty µL of 500 µM enterobactin in each experimental condition (with or without catalase) were added to three replicate wells on a 384-well plate, starting at time 0. Every 2 h, another row of 20 µL triplicates were added, until 6 h, and incubated at 37 °C in a Tecan Infinite M Nano spectrophotometer (Tecan Inc. Männedorf, Switzerland). At 6 h, fresh enterobactin 20 µL aliquots were added for the 0-time point and samples representing all time points were assayed using Arnow’s assay by adding 2 µL 5N HCl, 10 µL Arnow’s reagent, followed by 2 µL 10 N NaOH. Catecholate absorbance was measured with an Infinite M Nano spectrophotometer (Tecan Inc.) at 500 nm.

### Calmagite testing of pure enterobactin

Enterobactin is best known for its ability to bind the insoluble trivalent iron (Fe^3+^). In order to assess the ability of enterobactin purified from *R. mucilaginosa* to bind magnesium and zinc, the compleximetric dye calmagite was used in an assay format (40,41). Calmagite forms colored complexes with magnesium and zinc as well as other metals and was developed to quantify magnesium in biological samples (40). A chelating compound such as EDTA or enterobactin, is able to break this complex and elicit a color change from red to blue that can be measured spectrophotometrically. This assay was adapted to the analysis of enterobactin as follows: Calmagite dye (Acros Organics), was diluted in DI H_2_O to 0.4 mg/mL. A working solution was prepared consisting of a 25 µM metal ion solution (either MgSO_4_ or ZnSO_4_ diluted in DI H_2_O), and 0.05 mg/mL calmagite compleximetric dye diluted in 62.5 mM NH_4_Cl, pH10. One hundred and seventy-five µL were mixed with 25 µL of either a dilution series of EDTA (for quantification via a standard curve) or 25 µL of enterobactin sample at 896 µM and 100 µM (unknowns). The standard curve and unknown samples were all assayed in triplicate, and the disappearance of the red calmagite-metal complex due to EDTA or siderophore metal-binding competition was measured at 650 nm in a Tecan Infinite M Nano spectrophotometer set at room temperature. EDTA equivalents of enterobactin were calculated by extrapolating enterobactin absorbance values from 3^rd^ order polynomial line fitting of standard curve data (Fig. S7).

## Acknowledgements

Thanks to M. Donia, Princeton University, NJ, for sharing the *Actinomyces timonensis* DSM 23838 used in this work. Special thanks go out to M. Ghassemian, director of the University of California, San Diego Biomolecular and Proteomics Mass Spectrometry Facility, La Jolla, CA, for allowing use of the ABI 5600 mass spectrometer. Thanks to E, Glukhov at the University of California, Scripps Institution of Oceanography, La Jolla, CA for instrumentation support. This study was supported by NIH/NIDCR grant R00-DE024543 (A.E.).

## References

1. Lux T, Nuhn M, Hakenbeck R, Reichmann P. Diversity of bacteriocins and activity spectrum in Streptococcus pneumoniae. J Bacteriol. 2007;189(21):7741–51.

2. Nes IF, Diep DB, Holo H. Bacteriocin diversity in Streptococcus and Enterococcus. J Bacteriol. 2007;189(4):1189–98.

3. Tang X, Kudo Y, Baker JL, LaBonte S, Jordan PA, McKinnie SMK, et al. Cariogenic Streptococcus mutans Produces Tetramic Acid Strain-Specific Antibiotics That Impair Commensal Colonization. ACS Infect Dis. 2020;

4. Aleti G, Baker JL, Tang X, Alvarez R, Dinis M, Tran NC, et al. Identification of the Bacterial Biosynthetic Gene Clusters of the Oral Microbiome Illuminates the Unexplored Social Language of Bacteria during Health and Disease. 2019;10(2):1–19.

5. Donia MS, Cimermancic P, Schulze CJ, Wieland Brown LC, Martin J, Mitreva M, et al. A systematic analysis of biosynthetic gene clusters in the human microbiome reveals a common family of antibiotics. Cell. 2014;158(6):1402–14.

6. Donia MS, Fischbach MA. Small Molecules from the Human Microbiota. Science (80-). 2015;349(6246):139–48.

7. Qi B, Han M. Microbial Siderophore Enterobactin Promotes Mitochondrial Iron Uptake and Development of the Host via Interaction with ATP Synthase. Cell [Internet]. 2018;175(2):571-582.e11. Available from: https://doi.org/10.1016/j.cell.2018.07.032

8. Stubbendieck RM, May DS, Chevrette MG, Temkin MI, Wendt-Pienkowski E, Cagnazzo J, et al. Competition among nasal bacteria suggests a role for siderophore-mediated interactions in shaping the human nasal microbiota. Appl Environ Microbiol. 2019;85(10):1–17.

9. Ahmed E, Holmström SJM. Siderophores in environmental research: Roles and applications. Microb Biotechnol. 2014;7(3):196–208.

10. Achard MES, Chen KW, Sweet MJ, Watts RE, Schroder K, Schembri MA, et al. An antioxidant role for catecholate siderophores in Salmonella. Biochem J. 2013;454(3):543–9.

11. Peralta DR, Adler C, Corbalán NS, Paz García EC, Pomares MF, Vincent PA. Enterobactin as part of the oxidative stress response repertoire. PLoS One. 2016;11(6):1–15.

12. Adler C, Corbalan NS, Peralta DR, Pomares MF, D. Cristóbal RE, Vincent PA. The alternative role of enterobactin as an oxidative stress protector allows Escherichia coli colony development. PLoS One. 2014;9(1):1–10.

13. Bergan T, Kocus M. Stomatococcus mucilaginosus gen.nov., sp.nov., ep. rev., a member of the family Micrococcaceae. 1982;32(3):374–7.

14. Lim YW, Schmieder R, Haynes M, Furlan M, Matthews TD, Whiteson K, et al. Mechanistic Model of Rothia mucilaginosa Adaptation toward Persistence in the CF Lung, Based on a Genome Reconstructed from Metagenomic Data. PLoS One. 2013;8(5).

15. Baker JL, Morton JT, Dinis M, Alverez R, Bran NC, Knight R, et al. Deep metagenomics examines the oral microbiome during dental caries, 2 revealing novel taxa and co-occurrences with host molecules. 2019;0–1.

16. Iwasawa K, Suda W, Tsunoda T, Oikawa-kawamoto M. Dysbiosis of the salivary microbiota in pediatric-onset primary sclerosing cholangitis and its potential as a biomarker. Sci Rep [Internet]. 2018;(March):1–10. Available from: http://dx.doi.org/10.1038/s41598-018-23870-w

17. Perera M, Al-hebshi NN, Perera I, Ipe D, Ulett GC, Speicher DJ, et al. Inflammatory Bacteriome and Oral Squamous Cell Carcinoma. J Dent Res. 2018;97(6):725–32.

18. Lee SH, Lee Y, Park JS, Cho Y, Yoon H Il, Lee C, et al. Characterization of Microbiota in Bronchiectasis Patients with Different Disease Severities. J Clin Med. 2018;7(429):1–10.

19. Vieira AR, Hiller NL, Powell E, Kim LHJ, Spirk T, Modesto A, et al. Profiling microorganisms in whole saliva of children with and without dental caries. Clin Exp Dent Res. 2019;5(4):438–46.

20. Dashper SG, Mitchell HL, Lê Cao KA, Carpenter L, Gussy MG, Calache H, et al. Temporal development of the oral microbiome and prediction of early childhood caries. Sci Rep. 2019;9(1):1–12.

21. Maraki S, Papadakis IS. Rothia mucilaginosa pneumonia: A literature review. Infect Dis (Auckl). 2015;47(3):125–9.

22. Gao B, Gallagher T, Zhang Y, Elbadawi-Sidhu M, Lai Z, Fiehn O, et al. Tracking Polymicrobial Metabolism in Cystic Fibrosis Airways: Pseudomonas aeruginosa Metabolism and Physiology Are. mSphere. 2018;3(2):1–6.

23. McCormack MG, Smith AJ, Akram AN, Jackson M, Robertson D, Edwards G. Staphylococcus aureus and the oral cavity: An overlooked source of carriage and infection? Am J Infect Control [Internet]. 2015;43(1):35–7. Available from: http://dx.doi.org/10.1016/j.ajic.2014.09.015

24. Ohara-Nemoto Y, Haraga H, Kimura S, Nemoto TK. Occurrence of staphylococci in the oral cavities of healthy adults and nasal-oral trafficking of the bacteria. J Med Microbiol. 2008;57(1):95–9.

25. Smith AJ, Robertson D, Tang MK, Jackson MS, MacKenzie D, Bagg J. Staphylococcus aureus in the oral cavity: A three-year retrospective analysis of clinical laboratory data. Br Dent J. 2003;195(12):701–3.

26. Gevers D, Knight R, Petrosino JF, Huang K, McGuire AL, Birren BW, et al. The Human Microbiome Project: A Community Resource for the Healthy Human Microbiome. PLoS Biol. 2012;10(8):6–10.

27. Engen SA, Rørvik GH, Schreurs O, Blix IJS, Schenck K. The oral commensal Streptococcus mitis activates the aryl hydrocarbon receptor in human oral epithelial cells. Int J Oral Sci. 2017;9(3):145–50.

28. La Mantia I, Varricchio A, Ciprandi G. Bacteriotherapy with Streptococcus salivarius 24SMB and Streptococcus oralis 89a nasal spray for preventing recurrent acute otitis media in children: A real-life clinical experience. Int J Gen Med. 2017;10:171–5.

29. Regev-Yochay G, Trzciński K, Thompson CM, Malley R, Lipsitch M. Interference between Streptococcus pneumoniae and Staphylococcus aureus: In vitro hydrogen peroxide-mediated killing by Streptococcus pneumoniae. J Bacteriol. 2006;188(13):4996–5001.

30. Blin K, Shaw S, Steinke K, Villebro R, Ziemert N, Lee SY, et al. antiSMASH 5.0: updates to the secondary metabolite genome mining pipeline. Nucleic Acids Res. 2019;47(W1):W81–7.

31. Leong SA, Neilands J. Siderophore production by phytopathogenic microbial species. 1982;218(2):351–9.

32. Arnow EL. Colorimetric determination of the components of 3,4-dihyroxyphenylalaninetyrosine mixtures. J Biol Chem. 1937;531–7.

33. Nagaraja P, Vasantha RA, Sunitha KR. A sensitive and selective spectrophotometric estimation of catechol derivatives in pharmaceutical preparations. Talanta. 2001;55(6):1039–46.

34. Abdelmohsen UR, Grkovic T, Balasubramanian S, Kamel MS, Quinn RJ, Hentschel U. Elicitation of secondary metabolism in actinomycetes. Biotechnol Adv [Internet]. 2015;33(6):798–811. Available from: http://dx.doi.org/10.1016/j.biotechadv.2015.06.003

35. Kelley, Lawrence A., Mezulis, Stefans, Yates, Christopher M., Wass, Mark N., Sternberg MJ. The Phyre2 web portal for protein modeling, prediction and analysis. Nat Protoc [Internet]. 2015;10(6):845–58. Available from: http://dx.doi.org/10.1038/nprot.2015-053

36. Wang M, Carver JJ, Phelan V V., Sanchez LM., Garg N., Peng Y., et al. Sharing and community curation of mass spectrometry data with GNPS. Nat Biotechnol. 2017;34(8):828–37.

37. Liu GY, Essex A, Buchanan JT, Datta V, Hoffman HM, Bastian JF, et al. Staphylococcus aureus golden pigment impairs neutrophil killing and promotes virulence through its antioxidant activity. J Exp Med. 2005;202(2):209–15.

38. Lan L, Cheng A, Dunman PM, Missiakas D, He C. Golden pigment production and virulence gene expression are affected by metabolisms in Staphylococcus aureus. J Bacteriol. 2010;192(12):3068–77.

39. Sumioka R, Nakata M, Okahashi N, Li Y, Wada S, Yamaguchi M, et al. Streptococcus sanguinis induces neutrophil cell death by production of hydrogen peroxide. PLoS One. 2017;12(2):1–19.

40. Chauhan UPS, Sarkar BCRAY. Use of Calmagite for the Determination of Traces Magnesium in Biological Materials. 1969;32:70–80.

41. Kanadhia KC, Ramavataram Dvss, Nilakhe SPD, Patel S. A study of water hardness and the prevalence of hypomagnesaemia and hypocalcaemia in healthy subjects of Surat district (Gujarat). Magnes Res. 2014;27(4):165–74.

42. Recio E, Aparicio JF, Rumbero Á, Martín JF. Glycerol, ethylene glycol and propanediol elicit pimaricin biosynthesis in the PI-factor-defective strain Streptomyces natalensis npi287 and increase polyene production in several wild-type actinomycetes. Microbiology. 2006;152(10):3147–56.

43. Edlund A, Yang Y, Hall AP, Guo L, Lux R, He X, et al. An in vitro biofilm model system maintaining a highly reproducible species and metabolic diversity approaching that of the human oral microbiome. Microbiome. 2013;1(1):1.

44. Ebling K, Brent R. Media Preparation and Bacteriological Tools. Curr Protoc Mol Biol. 2002;Suplement:1–7.

45. Ziemert N, Podell S, Penn K, Badger JH, Allen E, Jensen PR. The natural product domain seeker NaPDoS: A phylogeny based bioinformatic tool to classify secondary metabolite gene diversity. PLoS One. 2012;7(3):1–9.

46. Kessner D, Chambers M, Burke R, Agus D, Mallick P. ProteoWizard: Open source software for rapid proteomics tools development. Bioinformatics. 2008;24(21):2534–6.

47. Wang M, Carver JJ, Phelan V V., Sanchez LM, Garg N, Peng Y, et al. Sharing and community curation of mass spectrometry data with Global Natural Products Social Molecular Networking. Nat Biotechnol. 2016;34(8):828–37.

48. Giner G, Smyth GK. Statmod: Probability calculations for the inverse Gaussian distribution. R J. 2016;8(1):339–51.

49. R Core Team. R: A Language and Environment for Statistical Computing [Internet]. 2019. Available from: https://www.r-project.org

50. Wickham H. ggplot2: Elegant Graphics for Data Analysis. New York: Springer-Verlag; 2009.

51. Weller H. countcolors: Locates and Counts Pixels Within Color Range(s) in Images. 2019.

52. Cacciatore S, Saccenti E, Piccioli M. Hypothesis: The Sound of the Individual Metabolic Phenotype? Acoustic Detection of NMR Experiments. Omi A J Integr Biol. 2015;19(3):147–56.

