## Supplemental Table 1 and Supplemental Figure legends for "Commensal oral *Rothia mucilaginosa* produces enterobactin – a metal chelating siderophore"

### Supplemental Material

**Table S1.** Experimental $^{13}C$ and $^{1}H$ chemical shifts (ppm) of enterobactin produced by *Rothia mucilaginosa* ATCC 25296 in DMSO-$d_6$. All chemical shifts in this work are identical to the already characterized enterobactin compound (40).

| Atom # | $^{13}C$ (ppm) | Type | $^{1}H$ (ppm) | Multiplicity | J (Hz) |
|---|---|---|---|---|---|
| Solvent: DMSO-d6 | | | | | |
| 1 | 169.0 | C | - | - | - |
| 2 | 51.1 | CH | 4.90 | m | - |
| 3 | 63.2 | $CH_2$ | 4.63 | t | 9.8 |
| | | | 4.40 | dd | 11.1,4.1 |
| 4 | 168.5 | C | - | - | - |
| 5 | 114.7 | C | - | - | - |
| 6 | 148.1 | C | - | - | - |
| 7 | 145.7 | C | - | - | - |
| 8 | 119.1 | CH | 6.96 | d | 8.2 |
| 9 | 118.3 | CH | 6.73 | t | 7.9 |
| 10 | 118.1 | CH | 7.34 | d | 8.2 |
| NH | - | - | 9.13 | br | - |
| 6-OH | - | - | 11.62 | br | - |
| 7-OH | - | - | 9.47 | br | - |

"m"=multiplet, "t"=triplet, "d"=doublet, "dd"=doublet of doublets, "br"=broad "-"= not applicable.

**FIG S1.** Overview of peptide sequence alignments showing the closest homologs to the biosynthetic gene clusters (BGCs) encoding: **A)** enterobactin produced by *Rothia mucilaginosa* ATCC 25296, and **B)** enterobactin produced by *Escherichia coli* K-12. Alignments were obtained using the antiSMASH v. 5.0 program (bacterial version). Adenylation domains in both pathways were predicted to select 2,3-dihydroxybenzoic acid (dhb) and serine (ser) as substrates, respectively. **C)** Close-up view of peptide alignment of *R. mucilaginosa* ATCC 25296 BGC and its closets homolog pathways (i.e. mirubactin [14%] and perquinoline A, B, C [15%], steffimycin D [5%]). **D)** Close-up view of peptide alignment of *E. coli* strain K-12 BGC to its closets homolog pathways (i.e. turnerbactin [13%], enterobactin [12%], streptobactin [23%] etc). The *R. mucilaginosa* BGC could not be aligned with the *E. coli* BGC due to non-existing peptide sequence homology in any of the genes except the NRPS genes which showed 41% homology (Fig. S4).

**FIG S2.** Catecholate siderophore encoding biosynthetic gene clusters identified by the antiSMASH software in genomes of: I) *Rothia mucilaginosa* ATCC 25296; II) *R. dentocariosa* M567; and III) *R. aeria* F0184. Predicted core biosynthetic genes, iron transporting genes, and genes encoding protein with species specific functions (A-F) are highlighted.

**FIG S3.** A) *Rothia mucilaginosa* ATCC 25296 (Rm) growth was established first on BHI agar (100mM sucrose) (required for growth of the challenging species *Actinomyces timonensis* DSM 23838 [At] during aerobic condition). At was spotted adjacent to Rm and its growth was inhibited. B) *Streptococcus salivarius* SHI-3 (Sal) shows a growth boost and forms growth on top of Rm when plated adjacent to Rm on M9 minimal agar media (100mM sucrose).

**FIG S4.** Results from peptide sequence alignment analysis of the NRPS gene in the *Rothia mucilaginosa* ATCC 25296 cat-sid BGC using the Phyre2 protein structure prediction tool showed 41% sequence homology to the EntE/EntB fusion protein harbored by the enterobactin BGC from *Escherichia coli* JM109.

**FIG S5.** A) 1H NMR spectrum of enterobactin purified from *R. mucilaginosa* ATCC 25296. B) 2D heteronuclear single quantum correlation (HSQC) NMR spectrum of enterobactin purified from *R. mucilaginosa* ATCC 25296. C) 2D heteronuclear multiple bond coherence (HMBC) NMR spectrum of enterobactin purified from *R. mucilaginosa*

ATCC 25296. D) 2D proton correlation spectroscopy (H COSY) NMR spectrum of enterobactin purified from *R. mucilaginosa* ATCC 25296.

**FIG S6.** *Rothia mucilaginosa* ATCC 25296 inhibits pigment production in *Staphylococcus aureus* enterotoxin H producing strain ATCC 51811, and MRSA strain TCH70 growing on M9 agar plates with no catalase added (100 mM glycerol).

**FIG S7.** Standard curves for the calmagite compleximetric assay for siderophore activity. A) 25 µM $MgSO_4$ complexed with calmagite. B) 25 µM $ZnSO_4$ complexed with calmagite. Both curves arise from dilutions of EDTA from 0-500 µM added to the calmagite-metal complex at pH 10 and a color change from red to blue monitored at 650 nm (37, 38). Enterobactin at 100 µM bound metal ions equal to 40 µM EDTA.
