## Supplementary figures and images for "Commensal oral *Rothia mucilaginosa* produces enterobactin – a metal chelating siderophore"

### Figure S1

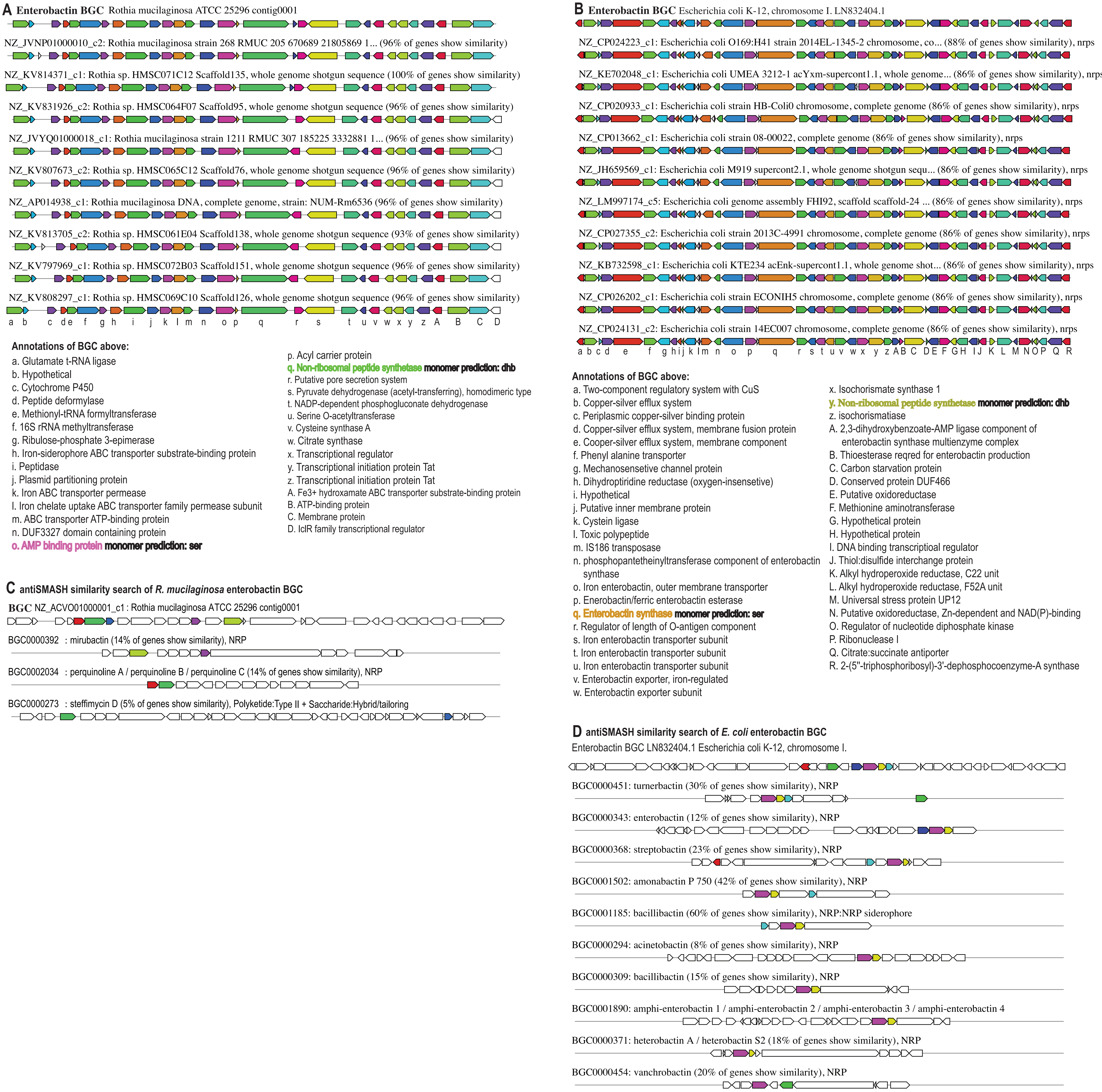

### Figure S2

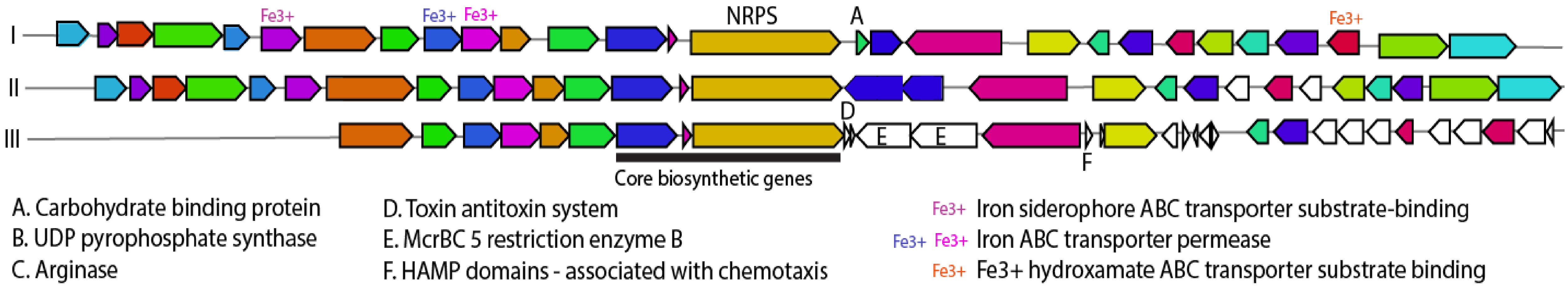

### Figure S3

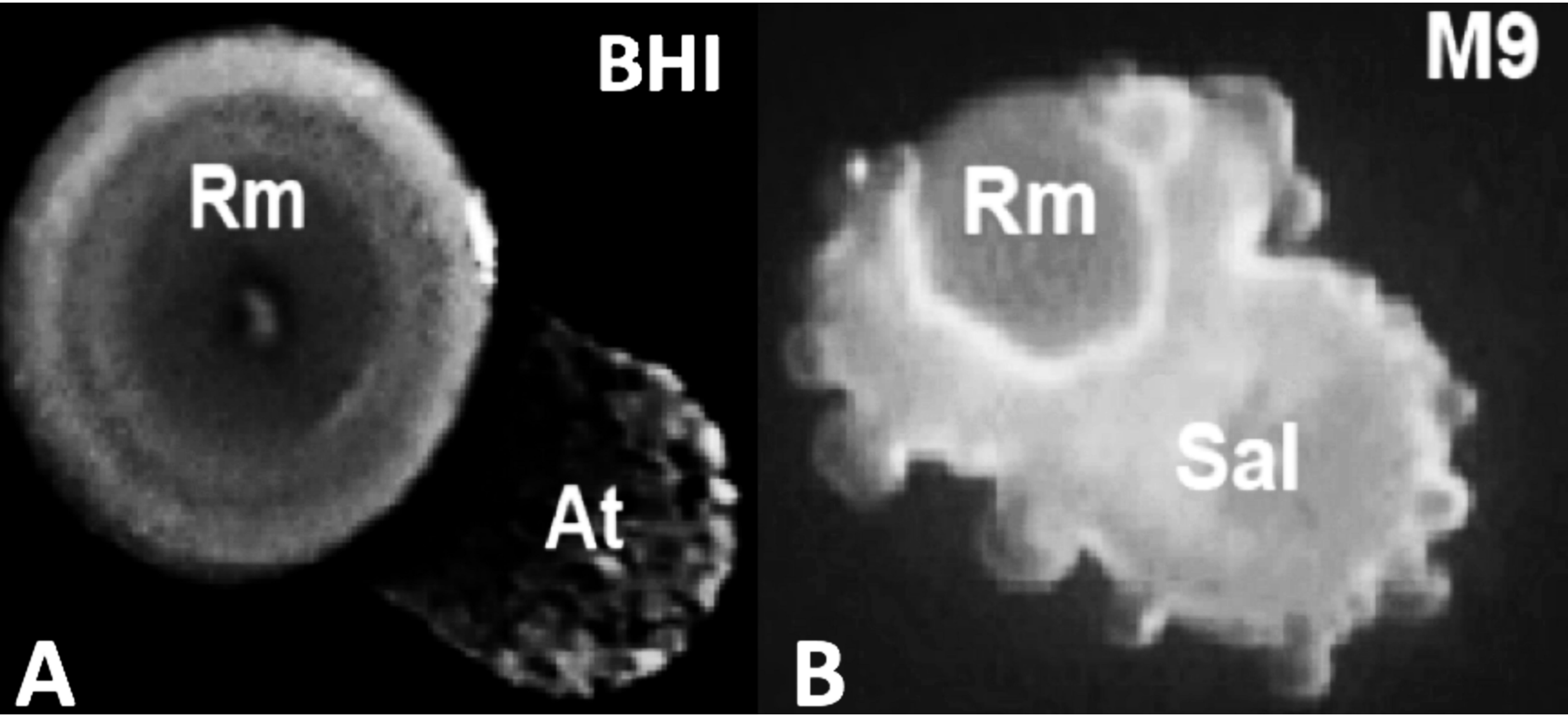

### Figure S4

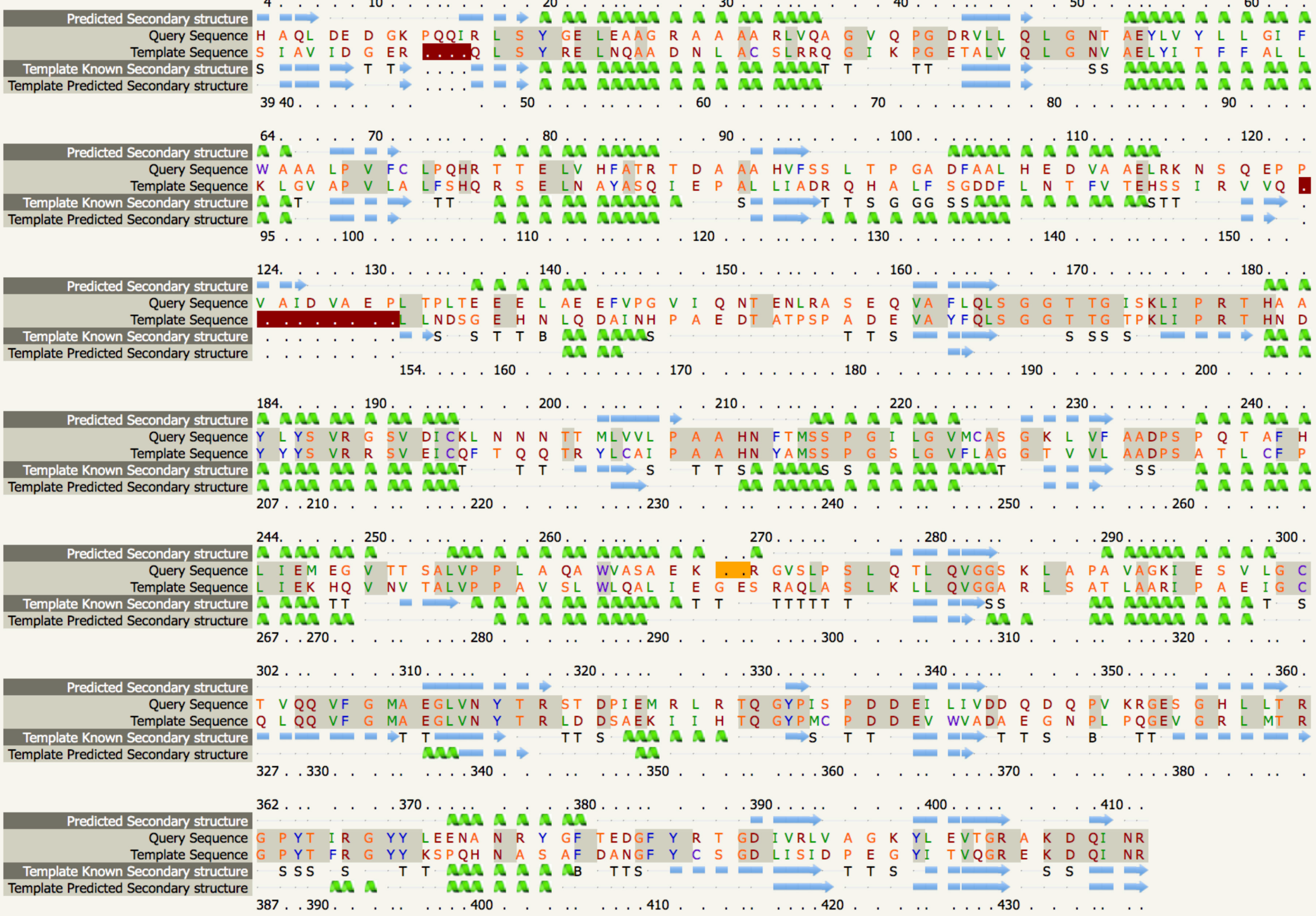

### Figure S5

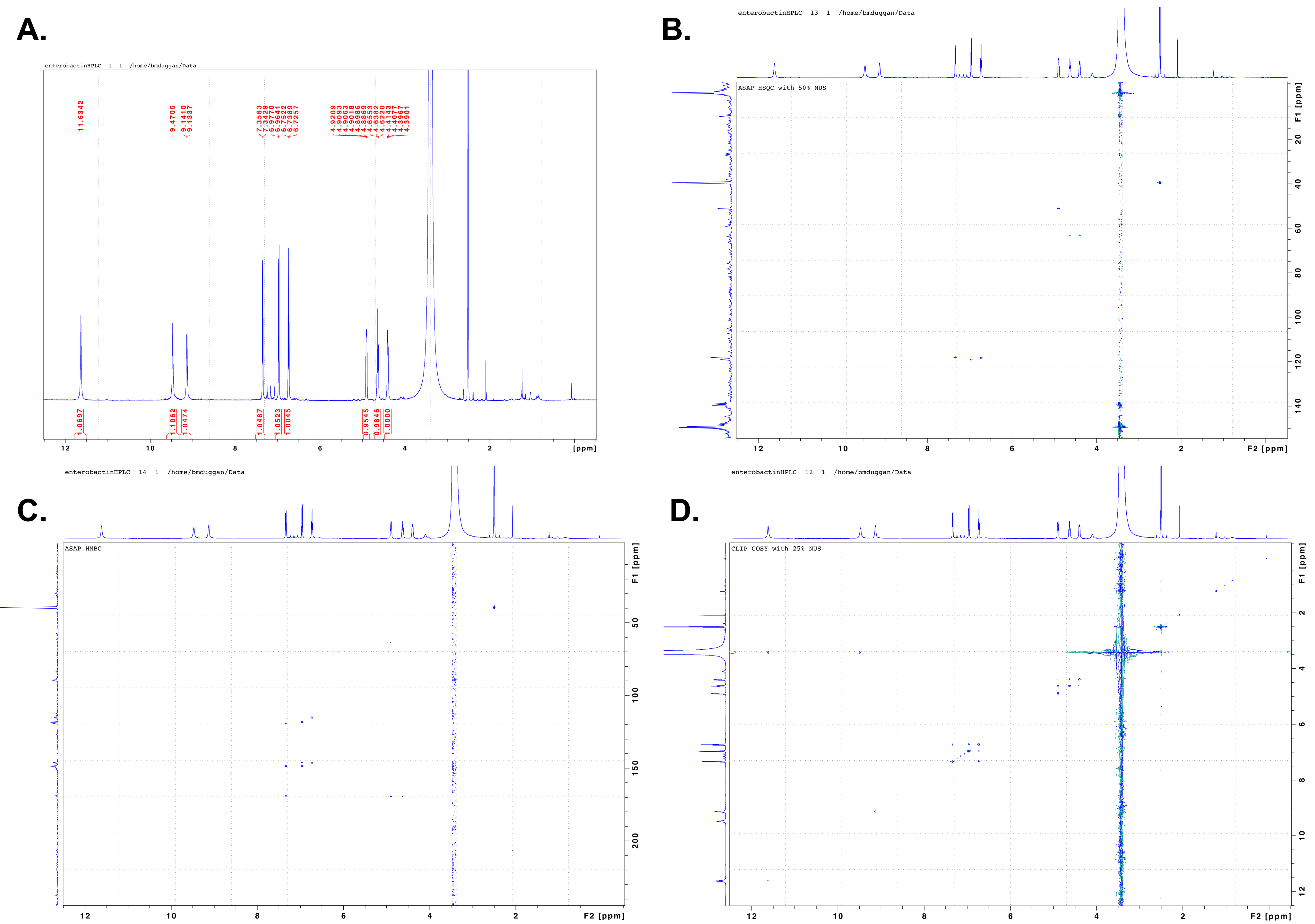

### Figure S7

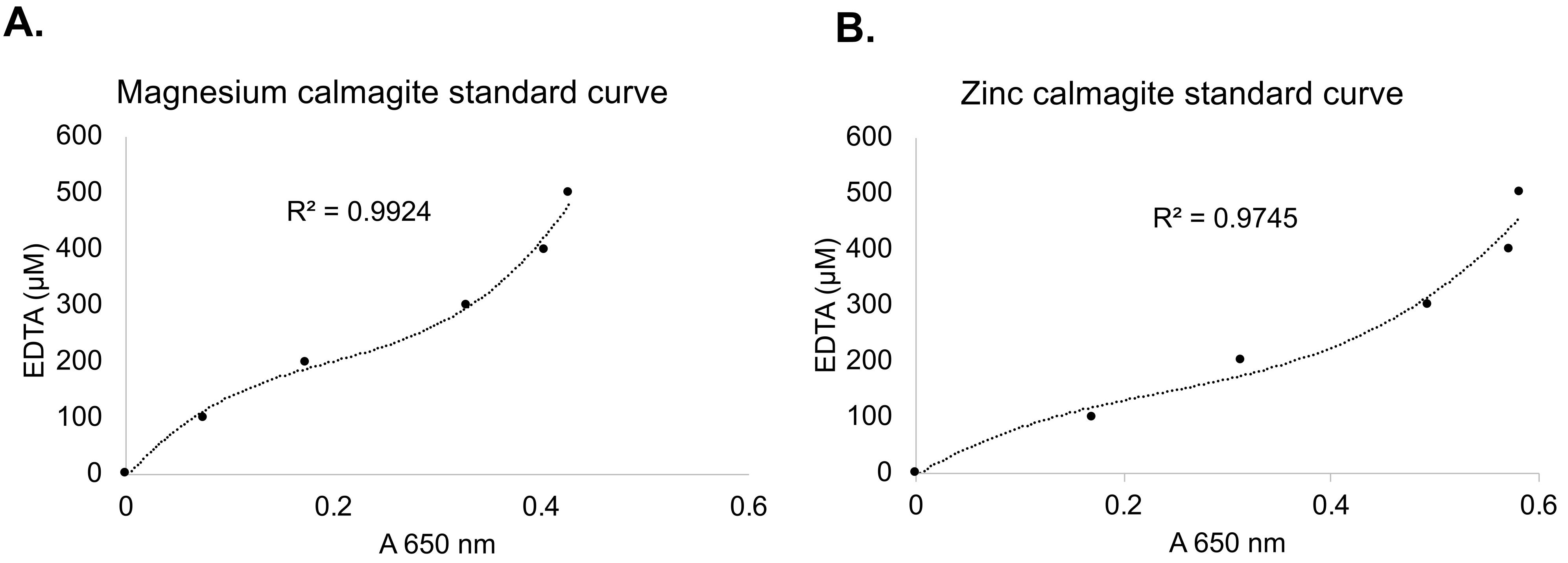
