## Supplementary material for "Commensal oral *Rothia mucilaginosa* produces enterobactin – a metal chelating siderophore": Figure S6

*R.muc* ATCC 25296
M9+glycerol

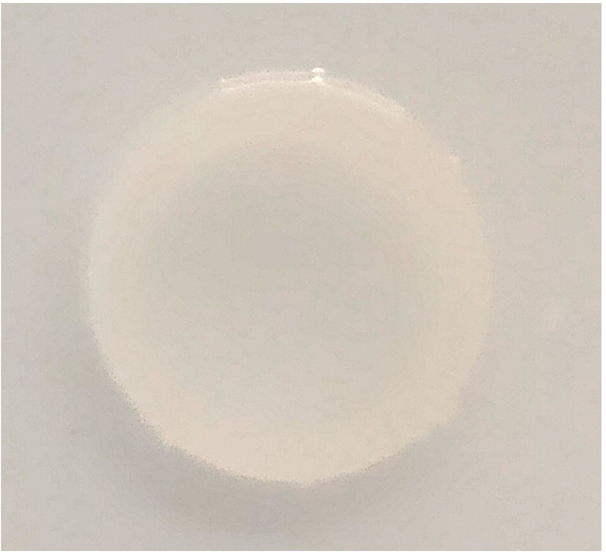

*S.aur* ATCC 51811
M9+glycerol

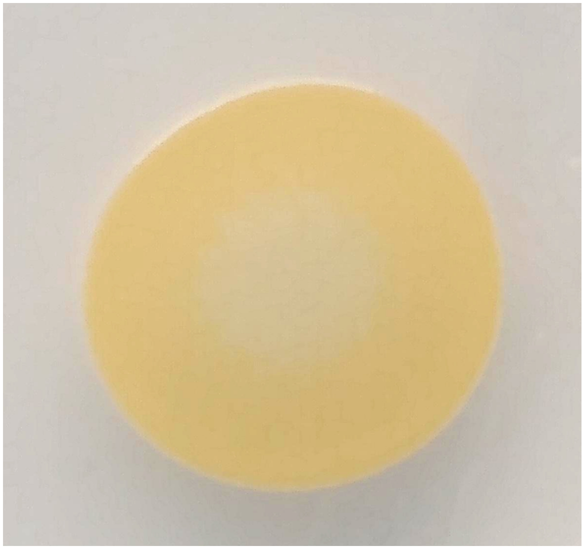

*R.muc*+*S.aur* ATCC 51811
M9+glycerol

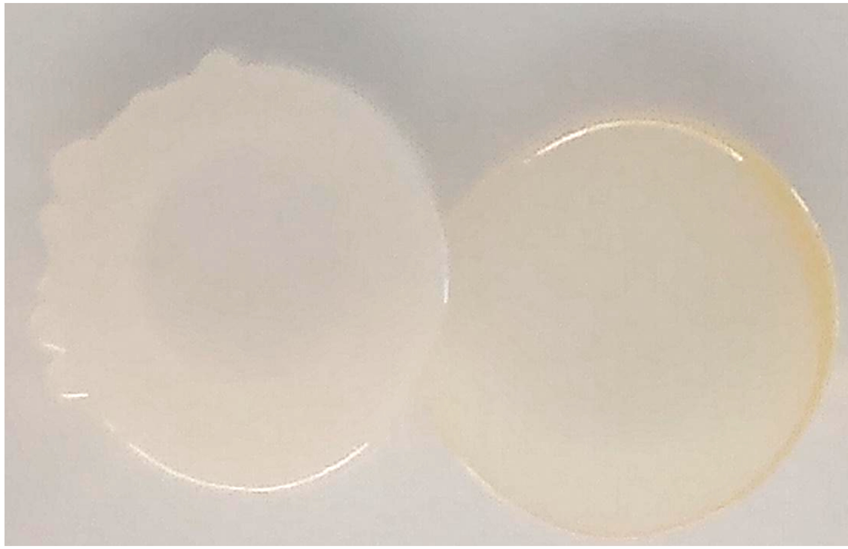

*S.aur* TCH 70 (MRSA)
M9+glycerol

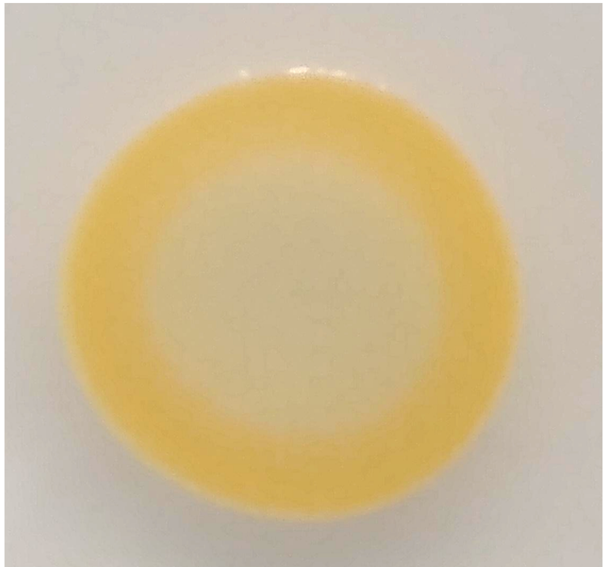

*R.muc*+*S.aur* TCH 70 (MRSA)
M9+glycerol

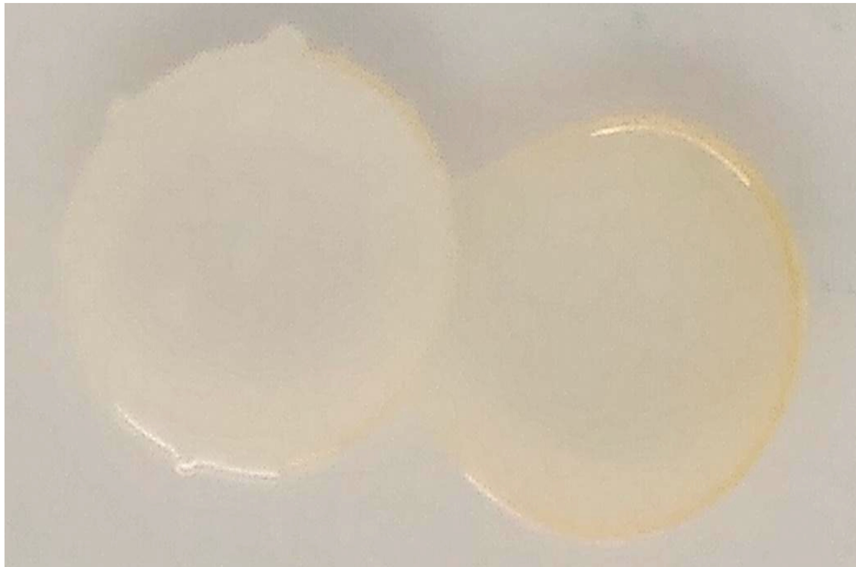
